# BrainAGE as a measure of maturation during early adolescence

**DOI:** 10.1101/2023.05.31.542949

**Authors:** Lucy B. Whitmore, Sara J. Weston, Kathryn L. Mills

## Abstract

The Brain-Age Gap Estimation (BrainAGE) is an important new tool that purports to evaluate brain maturity when used in adolescent populations. However, it is unclear whether BrainAGE tracks with other maturational metrics in adolescence. In the current study, we related BrainAGE to metrics of pubertal and cognitive development using both a previously validated model and a novel model trained specifically on an early adolescent population. The previously validated model was used to predict BrainAGE in two age bands, 9-11 and 10-13 years old, while the novel model was used with 9-11 year olds only. Across both models and age bands, an older BrainAGE was related to more advanced pubertal development. The relationship between BrainAGE and cognition was less clear, with conflicting relationships across the two models. Additionally, longitudinal analysis revealed moderate to high stability in BrainAGE across early adolescence. The results of the current study provide initial evidence that BrainAGE tracks with some metrics of maturation, including pubertal development. However, the conflicting results between BrainAGE and cognition lead us to question the utility of these models for non-biological processes.

## Introduction

Over the last 25 years, we have learned a considerable amount about the prolonged development of the human brain, due in part to advances in neuroimaging technologies and methods (Bethlehem et al., 2022; Mills et al., 2014; Raznahan et al., 2011; Wierenga et al., 2014). Brain development progresses differently across individuals (Mills et al., 2021), with different patterns and trajectories theorized to relate to outcomes such as psychopathology (Shaw et al., 2010). While longitudinal data is necessary to track individual trajectories, most neuroimaging studies that examine the brain across the lifespan are cross-sectional, and there has been an increasing desire to find ways to measure individual differences in brain maturation using methods that are compatible with cross-sectional data. The Brain-Age Gap Estimation (BrainAGE) has been proposed as one approach to assess the maturity of an individual’s brain, as this measure reflects the difference between an individual’s chronological age and their age as predicted by machine learning algorithms trained on neuroimaging data, often structural MRI (Brown et al., 2012; Franke et al. 2010). With this approach, an individual receives an estimated brain age that can differ from their chronological age.

### BrainAGE as a measure of aging and pathology

BrainAGE has most frequently been used in older adult populations, as it was originally proposed as a tool to research and assess risk for aging-related diseases (Franke et al., 2010). For older adults, differences between an individual’s estimated brain age and chronological age have been thought to reflect potential deviations from a normative aging trajectory related to various health concerns. This has been supported by previous work that has demonstrated having a BrainAGE that is older than one’s chronological age is related to metrics and outcomes of cognitive aging and decline, such as Alzheimer’s, in older adults (Franke & Gaser, 2012; 2019; Biondo et al., 2022). Additionally, across adulthood, having an older BrainAGE than one’s chronological age has also been associated with psychopathologies including depression and schizophrenia (Han et al., 2021; 2022; Schnack et al., 2016).

### BrainAGE in adolescence

Although BrainAGE is more commonly used in investigations of older adult populations, researchers are starting to use BrainAGE as a tool to advance current theories in developmental cognitive neuroscience. As BrainAGE was not originally developed as a measure of individual differences in adolescent brain maturation, the implications for interpreting BrainAGE are less clear in this population than in older populations where an older-appearing brain is more straightforwardly related to aging-related diseases. Conventionally, differences between BrainAGE and chronological age have been interpreted as reflecting accelerated or decelerated brain maturation in children and adolescents, though this has not been validated with longitudinal data (Franke et al., 2012; Lewis et al., 2018).

BrainAGE prediction based on structural MRI has been shown to accurately predict age in adolescents as measured by mean absolute errors in the one- to two-year range (Franke et al., 2012). Since the inception of using BrainAGE in adolescent populations, there have been efforts to link an adolescent’s BrainAGE to psychopathologies, which often emerge during this time. Recently, BrainAGE has shown promise as a potential biomarker of mental health issues in adolescent populations. Notably, having a BrainAGE that is older than one’s chronological age has been linked to major depression, functional impairment, and risk of psychosis (Drobonin et al., 2022; Chung et al., 2018; Copley et al., 2021). This is consistent with the idea that deviations from typical developmental trajectories play a role in the development of psychological disorders, as hypothesized by Shaw et al. (2010). Similarly, the Stress Acceleration Hypothesis (Callaghan & Tottenham, 2016) proposes that early life stress accelerates neural development, potentially resulting in short-term benefits but long-term vulnerability to psychopathology.

However, in recent years BrainAGE has also been theorized to reflect more than just accelerated or decelerated brain maturation (Vidal-Pineiro et al., 2021, Ball et al., 2021, Kelly et al., 2022). In adults, BrainAGE has been found to relate to birth weight and polygenic brain age scores (Videl-Pineiro et al., 2021). Additionally, adult BrainAGE varies over the course of the menstrual cycle (Franke et al., 2015). In children and adolescents, BrainAGE has been related to genetic factors (Brouwer et al., 2021). Despite this, the success of BrainAGE prediction in adolescent populations suggests its potential use as a clinically-relevant biomarker; however, given a lack of clarity around interpretations of adolescent BrainAGE, additional validation is needed before BrainAGE can be recommended for clinical and applied settings. In particular, it remains unclear as to which aspects of maturation are captured by this measure and which are not. A challenge to validation is the lack of conceptual clarity around the concept of “brain maturity” and what it means for a brain to be mature (Somerville, 2016).

The validation of any measure interpreted as reflecting a maturational process is an important step, and one that has been accomplished for developmental metrics such as the Tanner stages, also known as the Tanner Scale (Tanner, 1962). The Tanner stages, developed in the 1960s, describe and quantify physical changes associated with puberty. The stages were validated by observing adolescents and correlating physical changes with the corresponding stage, making it a widely accepted measure of physical maturation in clinical and research settings. In the case of adolescent BrainAGE, we would expect a similar relationship to be present, where BrainAGE is related to other maturational metrics that develop over adolescence. In the current study, we explore the relationship of BrainAGE to two domains of changes that are occurring in this time: pubertal and cognitive development.

### Brain maturation and puberty

Pubertal development is often measured in stages, such as Tanner stages or using the Pubertal Development Scale (PDS) (Tanner, 1962; Petersen et al., 1988). Tanner stages measure physical development, based on changes in pubic hair, genitalia, and breast growth (for females). Based on these metrics, Tanner Stage 1 represents pre-puberty, Stages 2-4 represent mid/intermediate puberty, and Stage 5 represents reproductive maturity. The Pubertal Development Scale examines similar, though not entirely overlapping, measures as Tanner stages and includes questions about characteristics such as body hair, growth in height, and other secondary sex characteristics (Petersen et al., 1988). Pubertal hormones such as testosterone or estradiol are also commonly examined in relation to pubertal development, but no current scale exists to assess pubertal development progress based on levels of certain hormones.

Independent of age, pubertal stage has been related to changes in brain structures, both cortical and subcortical. While cross-sectional relationships between subcortical volumes and pubertal development have found mixed results, longitudinal work has more consistently found a positive relationship between pubertal stage and amygdala and hippocampus volumes and a negative relationship between pubertal stage and nucleus accumbens, caudate, putamen, and globus pallidus volumes (Goddings et al., 2014; Herting and Sowell, 2017). Pubertal stage has also been found to be a better predictor of development than age in a number of subcortical regions, including the caudate, pallidum, and hippocampus (Wierenga et al., 2018).

A previous literature review found that both cross-sectional and longitudinal studies report widespread patterns of reductions in cortical gray matter in relation to pubertal development, both stage and timing (Vijayakumar et al., 2018). Notably, significant negative associations have been found between the pubertal hormone testosterone and cortical thinning in regions including the left posterior cingulate, precuneus, dlPFC and ACC in males and right somatosensory cortex in females (Nguyen, 2012). Additionally, change in Tanner stage has been related to rates of change in cortical surface area, where a main effect of stage was found for decreases in left precuneus surface area (Herting et al., 2015). More recent work has shown relationships between pubertal stage and increased cortical thinning in frontal and parietal cortices, as well as some temporal regions, independent of age (Vijayakumar et al., 2021).

In terms of pubertal development, recent work exists on the relationship between puberty and adolescent BrainAGE. Using a convolutional neural net based BrainAGE model trained on a lifespan population (5-93 years old), Holm et al., (2023) found a positive relationship between BrainAGE and parent-report pubertal development in a sample of 9-13 year olds from the ABCD study. Additionally, Holm and colleagues found a small association between the annualized rate of change in PDS scores and the annualized rate of change in BrainAGE, controlling for the annualized rate of change in chronological age.

Additionally, Dehestani et al., (2023a) found that earlier pubertal timing was associated with an older BrainAGE in a sample of 9-13 year old participants from the ABCD study. Pubertal timing was measured using “pubertal age”, a new measure that uses observed physical development (parent-report PDS) and pubertal hormones (testosterone and DHEA) to predict pubertal timing using a framework similar to that used for BrainAGE (Dehestani et al., 2023b). BrainAGE was predicted from 90 features of cortical volume, surface area, and thickness, as well as subcortical volume, using a support vector regression framework. The BrainAGE model was trained and tested on a sample of 9-13 year olds from the ABCD Study.

Both of the previous studies examined pubertal development in the ABCD Study cohort, but each captured unique aspects of the relationship between BrainAGE and puberty. Notably, the Holm analyses used a CNN-based lifespan model to examine pubertal stage, while the Dehestani model used was trained on explicit measurements of thickness, volume, and area and examined in relation to pubertal timing.

### Brain maturation and cognition

In addition to pubertal changes, adolescents undergo a number of cognitive changes, such as improved working memory performance and inhibitory control (Cromer et al. 2015; Davidson et al. 2006). Performance on the NIH Toolbox Cognition Battery, a commonly used measure of cognition, has been shown to improve cross-sectionally and longitudinally over early-to-mid adolescence (Anokhin et al., 2022). Additionally, working memory improvements in adolescence have been found to be associated with cortical volume reductions in bilateral prefrontal and posterior parietal regions and in regions around the central sulci (Tamnes et al, 2015).

### BrainAGE and cognition in adolescence

Past research on cognition and BrainAGE in adolescence has found mixed results, with some work indicating that a positive BrainAGE is related to faster processing speed (Erus et al., 2015) and others finding that a negative BrainAGE was associated with better cognitive performance (Lewis et al., 2018). Additionally, some studies have found very small or non-significant effects (Ball et al., 2017; 2021; Khudrakham et al., 2015, Kelly et al., 2022).

These studies exhibit a multitude of sources of variation, spanning from model features employed to train BrainAGE models, to participant age ranges, as well as the types of cognitive measures employed. In the studies described above, model features employed encompassed cortical thickness exclusively, T1 white/gray contrast, cortical thickness/volume/area, and subcortical volume, as well as the combination of gray matter, white matter, and ventricular measures. Parcellation methods varied significantly as well. Cognitive measurements employed in the studies were also highly varied, including IQ, measures of speed and accuracy on the Penn Computerized Neurocognitive Battery, and the NIH Toolbox Cognition Battery (Gur et al., 2012; Gershon et al., 2013). Although all of the studies incorporated adolescent participants, the age ranges varied significantly and the distribution of cognitive data across the age range was not always clearly indicated. Age ranges in certain studies were as broad as 8-22, 3-21, and 4.5-18.5 years, while Kelly et al. (2022) was restricted to cognition scores from participants who were 13 years old.

### Current Study

In the current study, we examine the relationship between BrainAGE and metrics of pubertal and cognitive maturation within two narrow age bands in early adolescence, 9-11 years and 10-13 years. For BrainAGE models trained on structural MRI data, such as the models in this paper, we expect that BrainAGE would track with maturational metrics that are reflected in changes to brain structure and change during these age ranges, including pubertal and cognitive development. Additionally, we examine the stability of BrainAGE in early adolescence in a longitudinal analysis.

The current study used three different BrainAGE models in order to provide additional confidence in our results by comparing models trained on both wide and narrow age ranges in order to balance specificity and replication. This strategy enabled us to balance providing the models with extensive data from our age range of interest, while not overly restricting our training data and possible predictions. In Study 1, we used an existing, previously validated BrainAGE model trained on a wide age range to predict BrainAGE in a sample of early adolescents (Drobonin et al. 2022).

In Study 2, we trained and tested two new BrainAGE model trained specifically on our age ranges of interest (9-11 years old and 10-13 years old). The development of BrainAGE models trained specifically on data from early adolescents is novel, and their usage reflects the unique structural changes occurring during this age range. The goal of the novel models is to capture the specific changes happening in these ranges, as opposed to the dynamic changes occurring over all of adolescence. While BrainAGE models exist that include adolescents, none sample early adolescence specifically, despite this being a unique time for structural brain development. The novel models are available for public use and can be found at https://github.com/LucyWhitmore/BrainAGE-Maturation.

Using predictions from all models, BrainAGE estimates were examined in association with youth-report pubertal development, parent-report pubertal development, and cognition scores.

As the goal of the study was to examine relationships between BrainAGE and other forms of maturation and determine whether these relationships replicate across models and age bands, the methods of Study 1 and Study 2 were kept as similar as possible. In particular, model training code, analysis procedures, and pubertal/cognitive measures were identical across Study 1 and Study 2. Differences in samples and procedures are described where present. In the following sections we discuss the studies separately, first detailing the procedures and results of Study 1, then the procedures and results of Study 2. Following Studies 1 and 2, we present longitudinal analyses of BrainAGE stability across both studies. Finally, we jointly discuss the implications, limitations, and future directions of the Studies 1 and 2.

#### Study 1

##### Participants

###### Model Training

The model used in Study 1 was a previously validated BrainAGE model, created for use in Drobonin et al. (2022). The model was trained on 1299 participants aged 9-19 years old, from a composite of six samples. Participants were members of the Autism Brain Imaging Data Exchange (ABIDE) Child Mind Institute: Healthy Brain Network (CMI), Consortium for Reliability and Validity, NIH MRI Study of Normal Brain Development, and Pediatric Imaging, Neurocognition, and Genetics (PING) cohorts. Distributions of age and gender per sample are available in Drobonin et al., (2022). Inclusion criteria included being within the age range of 9-19 years old and passing automated MRI quality control procedures. Additionally, only participants without psychiatric disorders and with an IQ over 75 were included in model training.

###### Model Prediction and Analyses

Adolescents included in the model predictions and analyses were participants in the Adolescent Brain Cognitive Development (ABCD) Study. The ABCD Study is a large longitudinal study that has recruited participants from 21 sites across the United States. Informed consent was obtained from all participants and their parents. For further details on recruitment procedures in the ABCD Study, see Garavan et al. (2018).

While the ABCD Annual Release 4.0 included both baseline data and data from the one- and two-year follow-ups, only baseline and the two-year follow-up included imaging data. Baseline imaging data were available for 11,878 participants between the ages of 9-11 years old. After excluding participants with missing or low-quality structural MRI data, 11,402 participants remained (*M* = 9.92 years old, *SD* = 0.63 years). Follow-up imaging data were available for 7,827 participants between the ages of 10 and 13. After excluding participants with missing or low-quality structural MRI data, 7,696 participants remained (*M* = 11.94 years old, *SD* = 0.65 years). Additional participant demographic details are available in *Table 1*.

**Table 1.** Participant demographics (ABCD).

|  |  | ***Baseline***<br>***(Model 1)*** | ***Follow-Up***<br>***(Model 1)*** | ***Baseline***<br>***(Model 2)*** | ***Follow-Up***<br>***(Model 2)*** |
| --- | --- | --- | --- | --- | --- |
| ***Sex*** | *Female* | 48%<br>($N=5471$) | 46.3% ($N = 3566$) | 48.4% ($N = 2759$) | 47.3% ($N = 1747$) |
| | *Male* | 52% ($N = 5931$) | 53.7% ($N = 4129$) | 51.6% ($N = 2943$) | 52.7% ($N = 1946$) |
| ***Race/***<br>***Ethnicity*** | *Non-Hispanic White* | 53.3%<br>($N=5962$) | 55.1%<br>($N=4242$) | 51.6%<br>($N=2946$) | 55.4%<br>($N=2045$) |
| | *Hispanic/Latino* | 20.5%<br>($N=2332$) | 19.3%<br>($N=1487$) | 21.5%<br>($N=1223$) | 19.7%<br>($N=729$) |
| | *Non-Hispanic Black* | 15.4%<br>($N=1754$) | 13.1%<br>($N=1010$) | 14.3%<br>($N=815$) | 12.9%<br>($N=476$) |
| | *Asian* | 2.1% ($N=244$) | 1.9% ($N = 145$) | 2.1% ($N = 122$) | 1.9% ($N=70$) |
| | *Native American/*<br>*Alaska Native* | 0.5% ($N = 56$) | 0.6% ($N = 44$) | 0.5% ($N = 30$) | 0.5% ($N=20$) |
| | *Multiracial* | 12% ($N = 1,370$) | 12.1% ($N = 931$) | 12.1% ($N = 688$) | 11.3%<br>($N=419$) |
| | *Additional Race* | 4.6% ($N = 528$) | 4.3% ($N = 335$) | 4.8% ($N = 277$) | 4.5%<br>($N=167$) |
| ***Household***<br>***Income*** | <\$5000 | 3.4 % ($N = 392$) | 2.5% ($N = 196$) | 3.3% ($N = 188$) | 2.6 % ($N=96$) |
| | \$5,000 - \$11,999 | 3.5% ($N = 398$) | 2.7% ($N = 211$) | 3.5% ($N = 199$) | 3% ($N=111$) |
| | \$12,000 - \$15,999 | 2.3% (N =265) | 1.8% (N =140) | 2.3% (N =130) | 1.9% (N=72) |
| | \$16,000 - \$24,999 | 4.4% (N =497) | 3.2% (N =246) | 4.2% (N =238) | 4% (N=148) |
| | \$25,000 - \$34,999 | 5.6% (N =633) | 5.2% (N =402) | 5.7% (N =324) | 5.6% (N=208) |
| | \$35,000 - \$49,999 | 7.8% (N =889) | 6.8% (N =524) | 8.3% (N =471) | 8.4% (N=309) |
| | \$50,000 - \$74,999 | 12.5% (N =1,429) | 12.7% (N =980) | 13.1% (N =747) | 14.3% (N=528) |
| | \$75,000 - \$99,999 | 13.3% (N =1,520) | 13.3% (N =1,026) | 12.8% (N =725) | 13.6% (N=503) |
| | \$100,000 - \$199,999 | 28.1 % (N =3,201) | 31.4% (N =2,413) | 27.6% (N =1,575) | 28.9% (N=1067) |
| | ≥ \$200,000 | 10.6% (N =1,208) | 12.49% (N =961) | 10.6% (N =603) | 9.9% (N=367) |
|  | Missing | 8.5% (N =970) | 7.7% (N =596) | 8.8% (N =501) | 7.7% (N=284) |
| **Model Training & Testing** | # Participants (Training) | 1,621 | 1,621 | 5,701 | 3,739 |
|  | Age Range (Training) | 9-19 y, m = 13.5 ± 3.04 | 9-19 y, m = 13.5 ± 3.04 | 8.9–11.08 y, m = 9.92 ± .63 | 10.58-13.67 y, m =11.95 ± .66 |
|  | # Participants (Testing) | 11,402 | 7,695 | 5,701 | 3,693 |
|  | Age Range (Testing) | 8.9–11.08 y, m = 9.92 ± .63 | 10.58-13.83 y, m = 11.94 ± .65 | 8.9–11.08 y, m = 9.92 ± .63 | 10.58-13.83 y, m = 11.94 ± .64 |

## Materials & Methods

### MRI Acquisition

For details on the acquisition of MRI images used to train the model used in Study 1, see Drobonin et al. (2022). For participants included in the analyses, detailed imaging procedures can be found in Casey et al. (2018) and Hagler et al. (2019). Harmonized protocols were used across 21 sites to collect imaging data using one of three 3T scanner platforms (Phillips, Siemens, or GE) and a 32-channel head coil; 3-dimensional T1-weighted scans were collected with 1-mm voxel resolution.

#### Model Training

As described in Drobonin et al., 2022, model training was conducted within the tidymodels (version 0.1.1) framework, using the XGBoost ML algorithm (version 1.0.0.2). Scan age was predicted from 189 features, including cortical and subcortical volume and area measurements.

### MRI Processing and Model Features

Model features included cortical gray matter volume and surface area measurements, as well as bilateral global and subcortical volume measurements. Cortical features were obtained using the Freesurfer default Desikan-Killiany atlas, while bilateral global and subcortical volume measures were obtained using the Freesurfer output. Data from the ABCD Study used for model testing were harmonized across sites using the longCombat package (Beer et al., 2020). The longCombat package was used to account for the multisite structure and longitudinal nature of the ABCD Study.

The Drobonin et al. (2022) model was originally trained on 189 structural features, but only 175 of these features were available in the ABCD dataset. Therefore, for the current study, the model was used to predict BrainAGE based on the 175 available features (Supplementary Table 1).

### BrainAGE Prediction

For the baseline sample, the Drobinin et al. (2022) model was used to predict BrainAGE for 11,402 adolescents (9-11 years old). For the two-year follow-up sample, the model was used to predict BrainAGE for 7,696 adolescents (10-13 years old).

### Bias Correction

Like other machine learning models, BrainAGE models can be susceptible to prediction bias towards the group mean (Le et al., 2018). In the case of BrainAGE models, this bias often takes the form of younger participants being more likely to be predicted as slightly older than their chronological age, while older participants are predicted to be slightly younger. To adjust for this age bias, we followed the bias correction procedures described in Smith et al. (2019). Briefly, the bias correction procedures require fitting a linear model to the validation set and extracting the intercept and slope. To obtain a corrected estimate, the intercept is subtracted from predicted age, and then value is then divided by the slope. The slope and intercept are generalizable to new data, allowing these coefficients to be used to correct test set bias (Peng et al., 2021). Additionally, age was included as a covariate in analyses to further correct for age-related bias.

### Behavioral & Survey Measures

#### Youth & Parent Report Pubertal Development

Both participants and their parents completed the Pubertal Development Scale (PDS; Petersen et al., 1988; Barch et al., 2018). The PDS is a five-item scale that measures pubertal stage. Of the five items, three are asked of every participant (related to growth in height, skin changes, and body hair growth), and two are dependent on the participants’ assigned sex at birth. The female version asks about the onset of menstruation and breast growth, while the male version asks about facial hair and vocal changes.

For each participant, mean scores were calculated for each of their self-report and parent-report scores (α = .53-.81, see Supplementary Table 2 for reliability estimates of the measures). Participants with incomplete data were excluded from the corresponding analyses.

#### Cognition

Cognition was measured using the NIH Toolbox Cognition Battery (Bleck et al., 2013; Gershon et al., 2013b; Hodes et al., 2013). The Toolbox uses seven cognitive tasks, which measure domains such as cognitive control, working memory, set shifting, and reading ability.

**Table 2.** NIH Toolbox Cognition Tasks.

|  |  |
| --- | --- |
| NIH Toolbox Flanker® | Cognitive Control/Attention |
| NIH Toolbox List Sorting Working Memory Test® | Working Memory; Categorization; Information Processing |
| NIH Toolbox Dimensional Change Card Sort® | Flexible thinking; concept formation; set shifting |
| NIH Toolbox Oral Reading Recognition Test® | Reading Ability; Language; Academic Achievement |
| NIH Toolbox Pattern Comparison Processing Speed® | Processing Speed; Information Processing |
| NIH Toolbox Picture Sequence Memory Test® | Visuospatial sequencing & memory |
| NIH Toolbox Picture Vocabulary Test® | Language; Verbal intellect |

*NIH Toolbox cognition tasks and their respective cognitive domains.*

Summary scores were provided in the ABCD 4.0 data release for each participant based on their performance on the seven tasks, listed in Table 1 and described in detail in Luciana et al. (2018). Summary scores are presented as the participants’ performance compared to those in the NIH Toolbox normative sample, which includes nationally representative data from participants who are between 3-85 years old. Complete cognition summary scores were only available at baseline. Distributions of all maturational metrics are available in Table 3.

**Table 3.** Descriptive statistics.

|  | **BrainAGE - Corrected** |  | **Youth-Report Puberty** |  | **Parent-Report Puberty** |  | **Cognition** |  |
| --- | --- | --- | --- | --- | --- | --- | --- | --- |
| Study | Mean ± SD | Range | Mean ± SD | Range | Mean ± SD | Range | Mean ± SD | Range |
| Study 1, Baseline | 0.68 ± 1.75 | -4.97 - 8.52 | 1.71 ± 0.47 | 1.0 - 4.0 | 1.60 ± 0.48 | 1.0 - 4.0 | 86.34 ± 9.10 | 44 - 117 |
| Study 1, Follow-Up | 0.19 ± 1.98 | -5.73 - 7.90 | 2.07 ± 0.65 | 1.0 - 4.0 | 2.12 ± 0.70 | 1.0 - 4.0 | -- | -- |
| Study 2, Baseline | -0.05 ± 2.19 | -7.71 - 7.36 | 1.72 ± 0.48 | 1.0 - 4.0 | 1.60 ± 0.48 | 1.0 - 4.0 | 86.31 ± 9.15 | 44 - 115 |
| Study 2, Follow-Up | 0.01 ± 2.80 | -9.68 - 9.84 | 2.08 ± 0.65 | 1.0 - 4.0 | 2.12 ± 0.71 | 1.0 - 4.0 | -- | -- |

### Statistical Analyses

Statistical analyses were performed in RStudio (R version 4.1.1, RStudio version 2021.09.0). Associations with maturational metrics were analyzed with linear models, using the *lm()* function. Separate models were created for the relationship between BrainAGE and each maturational metric, where BrainAGE was regressed on each metric of puberty and cognition. Age was included as a covariate to reduce age-related bias in BrainAGE estimates. We report unstandardized effects and standard errors in-text and in Table 4. Standardized effects are available in Table 5. Full regression models are available in Supplementary Tables 3-11.

**Table 4.** Summary of results, unstandardized.

|  | Youth-Report Puberty |  |  | Parent-Report Puberty |  |  | Cognition |  |  |
| --- | --- | --- | --- | --- | --- | --- | --- | --- | --- |
|  | beta | SE | p | beta | SE | p | beta | SE | p |
| Study 1, Baseline | 0.25 | 0.05 | <.001 | 0.24 | 0.04 | <.001 | -0.003 | 0.002 | <.001 |
| Study 1, Follow-Up | 0.50 | 0.04 | <.001 | 0.58 | 0.03 | <.001 | -- | -- | -- |
| Study 2, Baseline | 0.31 | 0.09 | <.001 | 0.19 | 0.06 | <.001 | 0.002 | 0.003 | .005 |
| Study 2, Follow-Up | 0.39 | 0.08 | <.001 | 0.42 | 0.07 | <.001 | - | - | - |

**Table 5.**
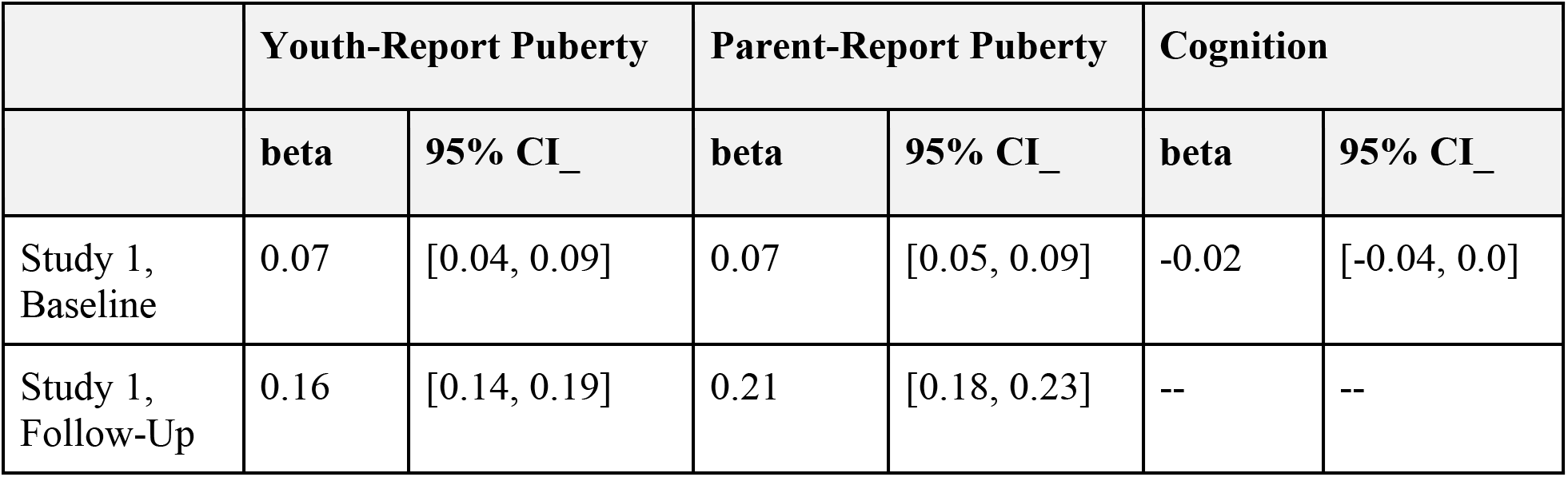
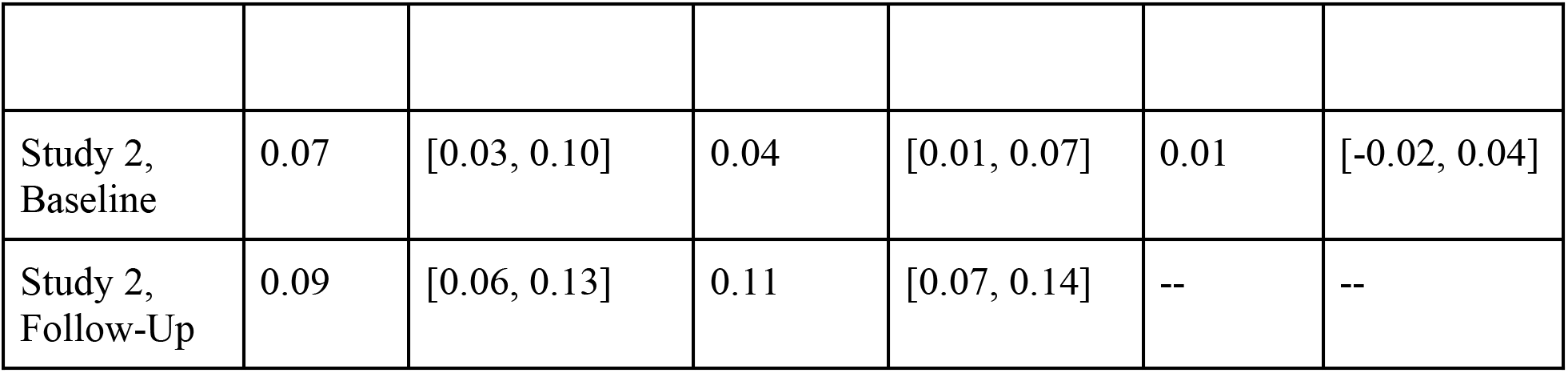
Summary of results, standardized.

|  | Youth-Report Puberty |  | Parent-Report Puberty |  | Cognition |  |
| --- | --- | --- | --- | --- | --- | --- |
|  | beta | 95% CI_ | beta | 95% CI_ | beta | 95% CI_ |
| Study 1, Baseline | 0.07 | [0.04, 0.09] | 0.07 | [0.05, 0.09] | -0.02 | [-0.04, 0.0] |
| Study 1, Follow-Up | 0.16 | [0.14, 0.19] | 0.21 | [0.18, 0.23] | -- | -- |
| Study 2, Baseline | 0.07 | [0.03, 0.10] | 0.04 | [0.01, 0.07] | 0.01 | [-0.02, 0.04] |
| Study 2, Follow-Up | 0.09 | [0.06, 0.13] | 0.11 | [0.07, 0.14] | -- | -- |

## Results

### BrainAGE Model Performance

In the original validation of the model, the model performed with a corrected mean absolute error (MAE) of 1.98 (Drobonin et al., 2022). This error is in line with prior adolescent BrainAGE models, which have performed with MAEs frequently in the 1-2 year range (Franke & Gaser, 2019). The existing model performed with a MAE of 2.32 before correction for age bias, and a corrected MAE of 1.45 on the ABCD data (baseline) and an uncorrected MAE of 1.3 (corrected MAE 1.58) at follow-up. Model performance is shown in *Figure 1*. A slight age bias was observed, as is common in these types of models, and was discussed previously. BrainAGE (predicted age subtracted from chronological age) had a correlation of −0.1 with chronological age at baseline, and −0.01 at follow-up.

**Figure 1.**
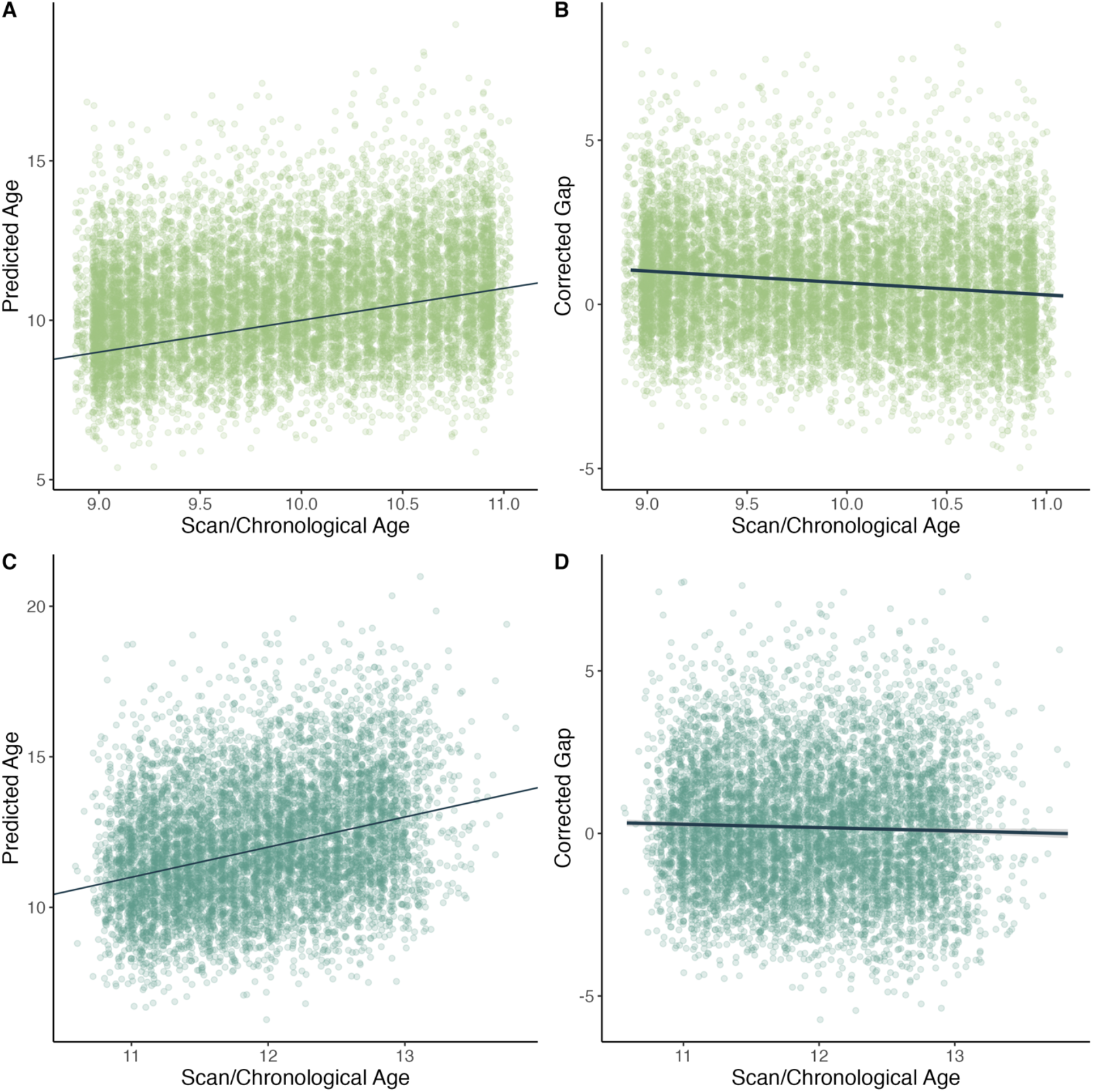
Age versus model predictions for the previously validated model. A) Scan/chronological age versus model-predicted age for the baseline sample. The line represents an exact relationship between chronological age and predicted age. B) Scan/chronological age versus the age-corrected brain-age gap, for the baseline sample. The regression line shows a slight age-related bias, with younger participants predicted to be slightly older than their chronological age. C) Similar to a), scan/chronological age versus predicted age for the follow-up sample. D) Similar to b) scan/chronological age versus the age-corrected brain-age gap for the follow-up sample. Again, the regression line shows a slight amount of age-bias, but less than at baseline.

### BrainAGE and Maturational Metrics

In the baseline wave (9-11 years old), a higher/more positive BrainAGE was related to more advanced youth- and parent-report pubertal development (Table 4). For youth-report pubertal development, (*b_youth_* = 0.25, *se* = 0.05*, p* < .001) *(Figure 2a)*. For parent-report pubertal development, (*b_parent_* = 0.24, *se* = 0.04*p* < .001) *(Figure 2b)*. Additionally, a higher/more positive BrainAGE was also related to lower cognition scores (*b_cog_* = −0.003, *se* = 0.002, *p* < .001 in both samples) *(Figure 2c)*.

**Figure 2.**
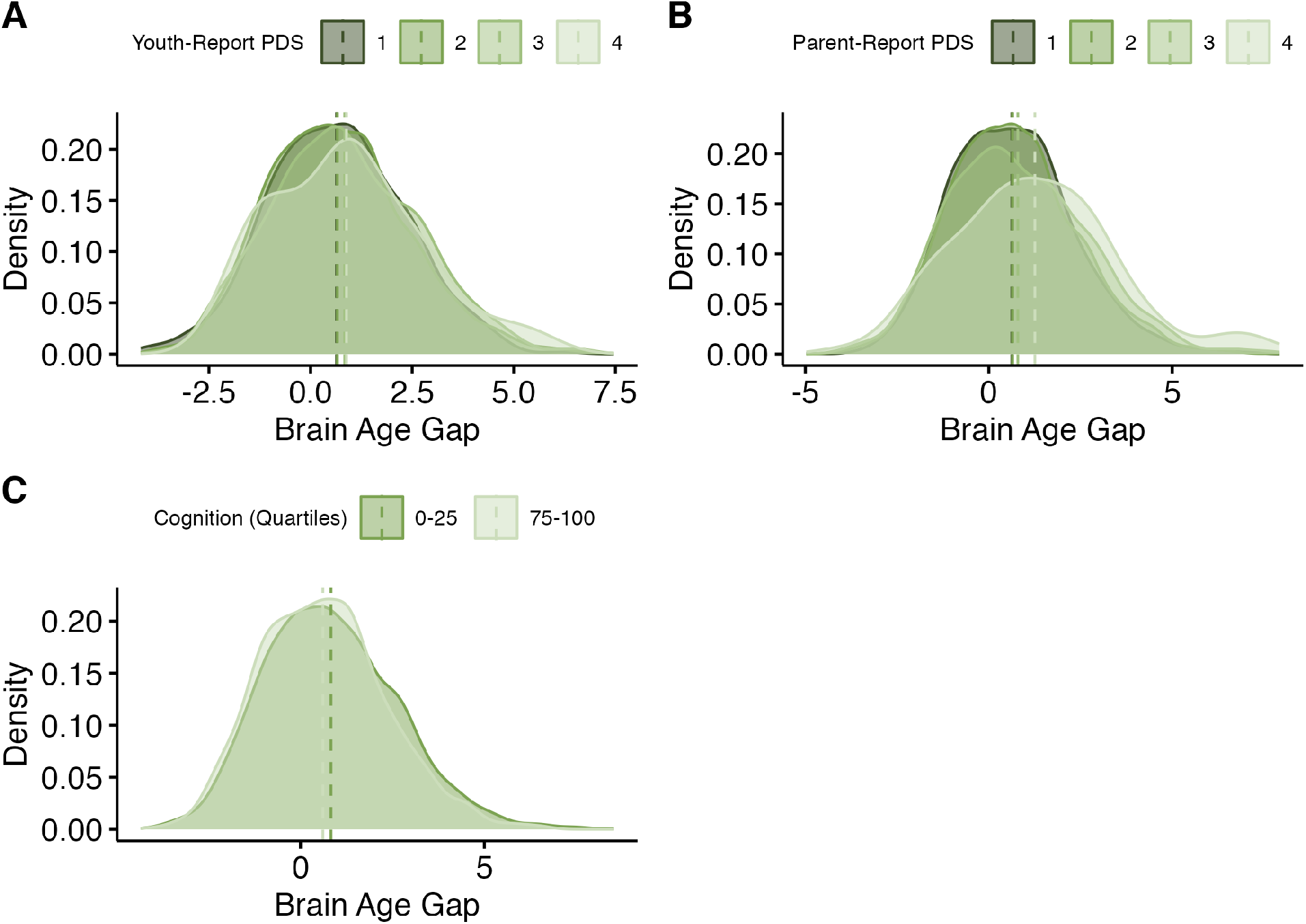
Results - Previously validated model, baseline sample. a) Distributions of brain-age gaps, grouped by pubertal stage (youth-report). Means for each group are represented with vertical lines. b) Distributions of brain-age gaps, grouped by pubertal stage (parent-report). Means for each group are represented with vertical lines. c) Distributions of brain-age gaps, grouped by quartile of cognition scores, compared to others in the sample. Only the highest and lowest quartiles are shown. Means for each group are represented with vertical lines.

In the two-year follow-up, a higher/more positive BrainAGE was related to more advanced youth- and parent-report pubertal development. For youth-report pubertal development, (*b_youth_*= 0.50, *se* = 0.04, *p* < .001) *(Figure 3a)*. For parent-report pubertal development, (*b_parent_*=0.58, *se* = 0.03, *p* < .001 in both samples) *(Figure 3b)*. As discussed above, composite cognition scores were not available at the two-year follow-up.

**Figure 3.**
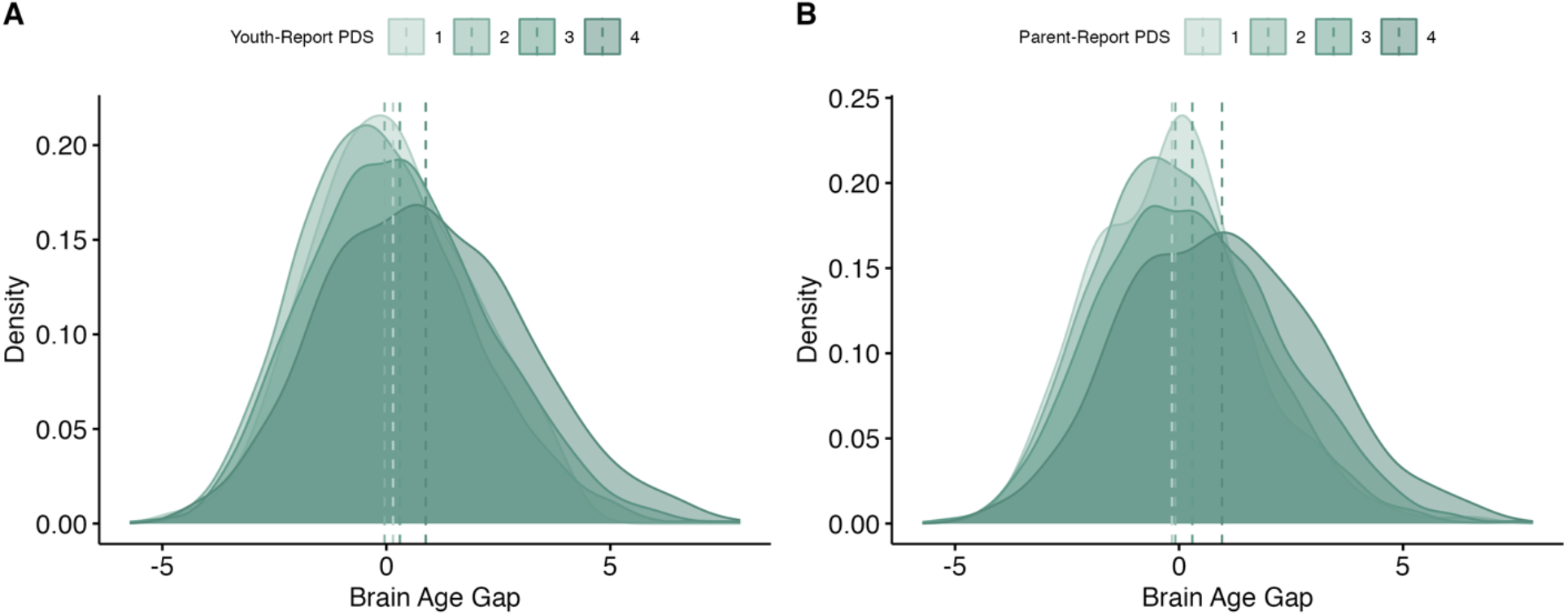
Results - Previously validated model, follow-up sample. a) Distributions of brain-age gaps, grouped by pubertal stage (youth-report). Means for each group are represented with vertical lines. b) Distributions of brain-age gaps, grouped by pubertal stage (parent-report). Means for each group are represented with vertical lines.

#### Study 2

##### Participants

###### Model Training and Analyses

Adolescents included in the model training, testing, and analyses were participants in the Adolescent Brain Cognitive Development (ABCD) Study, described in Study 1. Data included in Study 2 - Baseline were from the baseline sample (9-11 years old) and are identical to the baseline sample described in Study 1. Data included in Study 2 – Follow-Up were from the two-year follow-up (10-13 years old) and are identical to the follow-up sample described in Study 1.

###### MRI Acquisition

Detailed imaging procedures can be found in Casey et al. (2018) and Hagler et al. (2019). Harmonized protocols were used across 21 sites to collect imaging data using one of three 3T scanner platforms (Phllips, Siemens, or GE) and a 32-channel head coil. 3-dimensional T1-weighted scans were collected with 1-mm voxel resolution.

## Materials & Methods

### MRI Processing and Model Features

The models were trained on and used to predict BrainAGE from the same 175 structural features used in the prediction for Study 1. A complete list of the features used in the training and prediction of Study 2 is available in *Supplementary Table 1*. As in Study 1, the longCombat package was used to harmonize multisite data.

### BrainAGE Prediction

For both Study 2 models, available data were split into training, validation, and test sets. For the baseline model, half the sample (5,701 participants) were reserved for the test set. Of the remaining 5,701 participants, eighty percent (*N* = 4,559) were used for model training, and twenty percent (*N* = 1,142) were reserved for model validation. For Study 2 – Follow-Up, approximately half the sample (3,693 participants) were reserved for the test set. The test set was structured to maximize the number of participants included from the Study 2 baseline test set, enabling further longitudinal analyses. Of the remaining 3,739 participants, eighty percent (*N* = 2989) were used for model training, and twenty percent (*N* = 750) were reserved for model validation.

Model training followed the same procedures as the Study 1 model, originally described in Drobonin et al. (2022). Ten-fold cross-validation was repeated 10 times, stratified by scan age. Model training was performed using tidymodels (version 0.2.0; Kuhn & Wickham, 2020), with the XGBoost ML algorithm (version 1.5.2.1; Chen & Guestrin, 2016). Scan age was predicted using the cortical and subcortical features in Supplementary Table 1. The best model was the one with the smallest MAE (mean absolute error), which was expressed in years. Following model validation, the brain age gap was calculated in the hold-out sample by subtracting the actual scan age/chronological age from the predicted age. Descriptive statistics for BrainAGE predictions and maturational metrics are available in Table 3.

## Results

### BrainAGE Model Performance

#### Model 2 - Baseline

The best baseline model performed with a MAE of 0.14 in cross-validation. In our hold-out validation sample, the model performed with a MAE of 0.49. Finally, in the sample reserved for analyses, the model performed with a MAE of 0.49 (uncorrected for age bias) and 0.86 (corrected). Model performance is shown in *Figure 4a,b*. Model performance was improved over existing models, including the model used in Study 1. The model showed a slight age bias, again due to the effect of regression to the mean. Corrected BrainAGE had a correlation of −0.20 with chronological age (Figure 4b).

**Figure 4.**
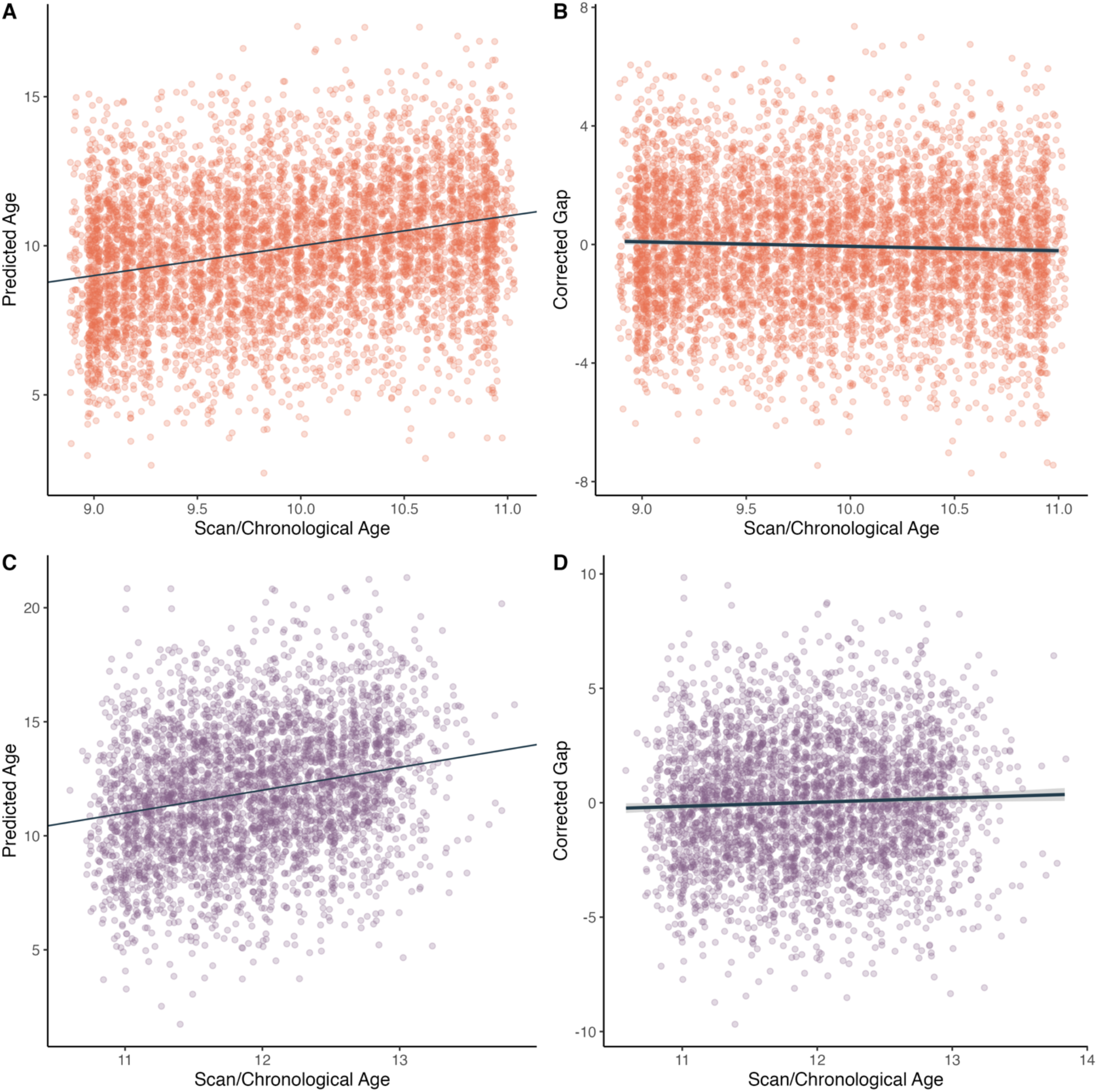
Age versus model predictions for the novel models. A) Scan/chronological age versus model-predicted age for the baseline sample. The line represents an exact relationship between chronological age and predicted age. B) Scan/chronological age versus the age-corrected brain-age gap estimate (BrainAGE), for the baseline sample. The regression line shows a slight age-related bias, with younger participants predicted to be slightly older than their chronological age. C) Similar to a), scan/chronological age versus predicted age for the follow-up sample. D) Similar to b) scan/chronological age versus the age-corrected brain-age gap for the follow-up sample. Again, the regression line shows a slight amount of age-bias.

##### Model 2 – Follow-Up

The best baseline model performed with a MAE of 0.53 in cross-validation. In our hold-out validation sample, the model performed with a MAE of 0.53. Finally, in the sample reserved for analyses, the model performed with a MAE of 0.52 (uncorrected for age bias) and 2.2 (corrected). Model performance is shown in *Figure 4c,d*. As with the baseline model, the novel follow-up model performed with similar accuracy to existing adolescent models. The model showed a slight age bias, again due to the effect of regression to the mean. Corrected BrainAGE had a correlation of 0.04 with chronological age (Figure 4d).

### Neuroanatomical Contributions

Top contributors to the models were determined using the vip R package (version 0.4.1), and visualized with the ggseg R package (version 1.6.5; Mowinckel & Vidal-Piñeiro, 2020). For models built using xgboost, the vip package determines variable importance by ranking the absolute magnitude of the estimated coefficients.

#### Baseline Model

The top contributors to the baseline model included the volume of the brainstem, mid-posterior corpus callosum, left posterior cingulate, right cuneus, left postcentral gyrus, and right isthmus cingulate.

*Figure 5* shows the cortical and subcortical features with the greatest importance to the model prediction.

**Figure 5.**
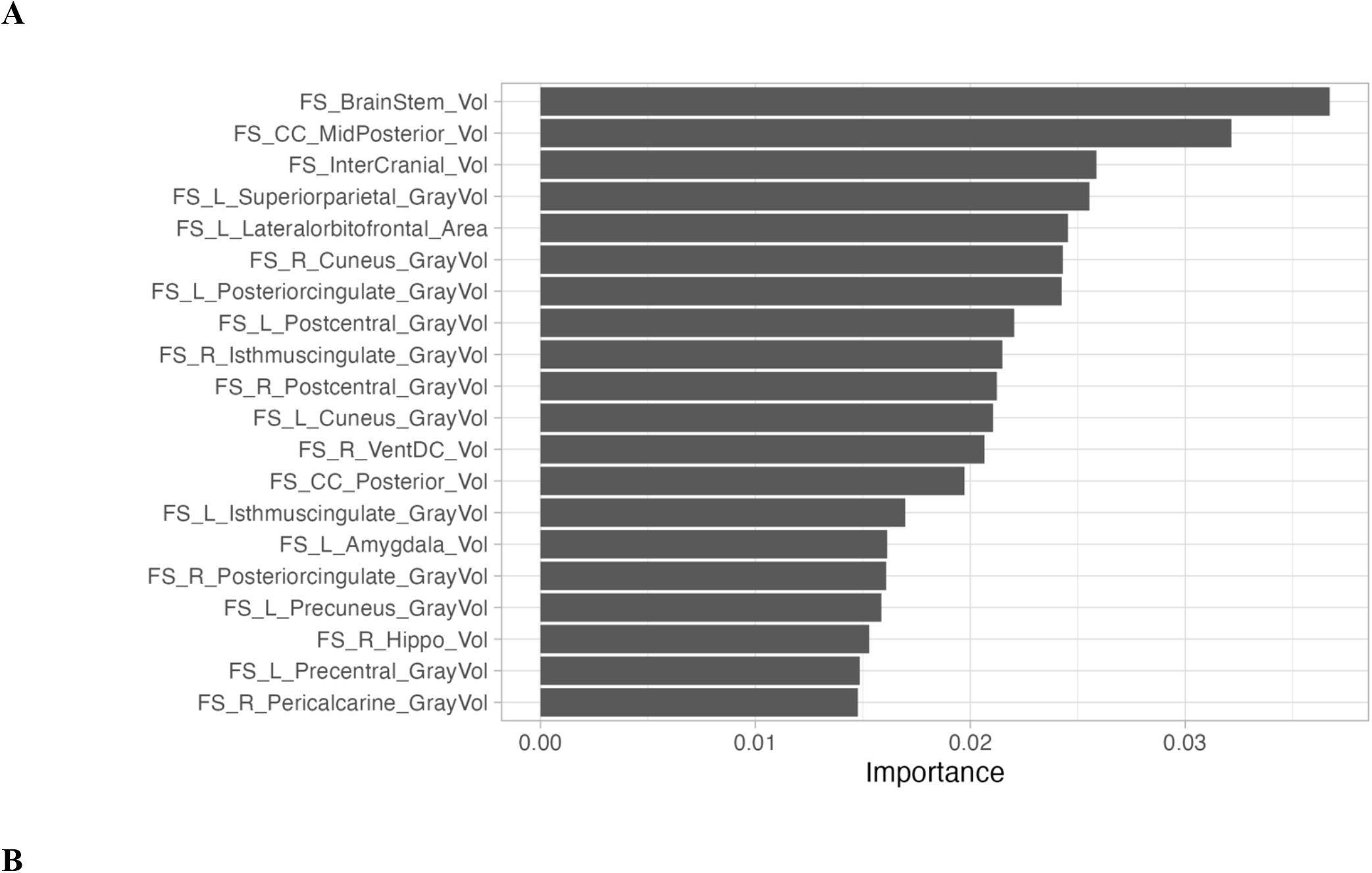
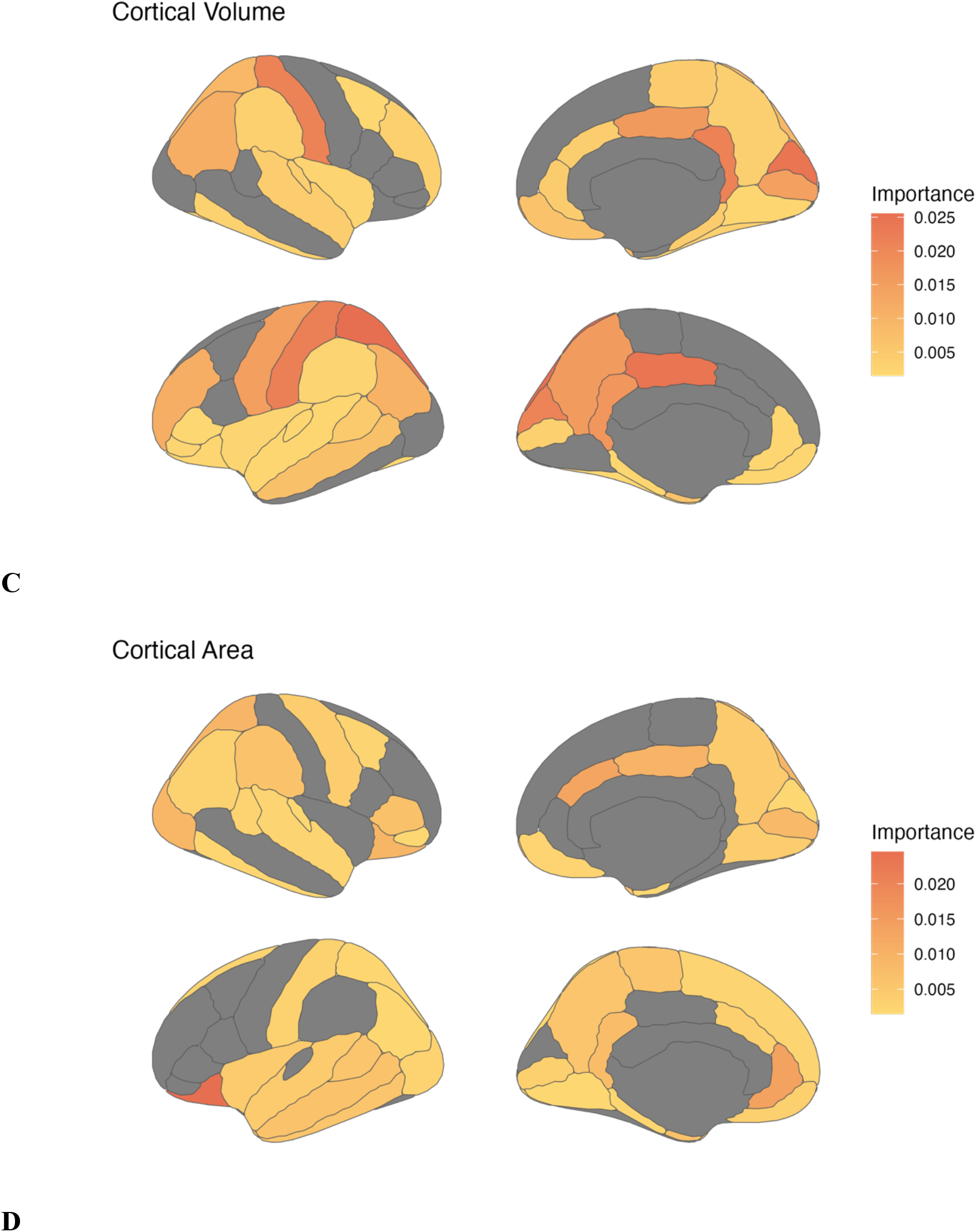
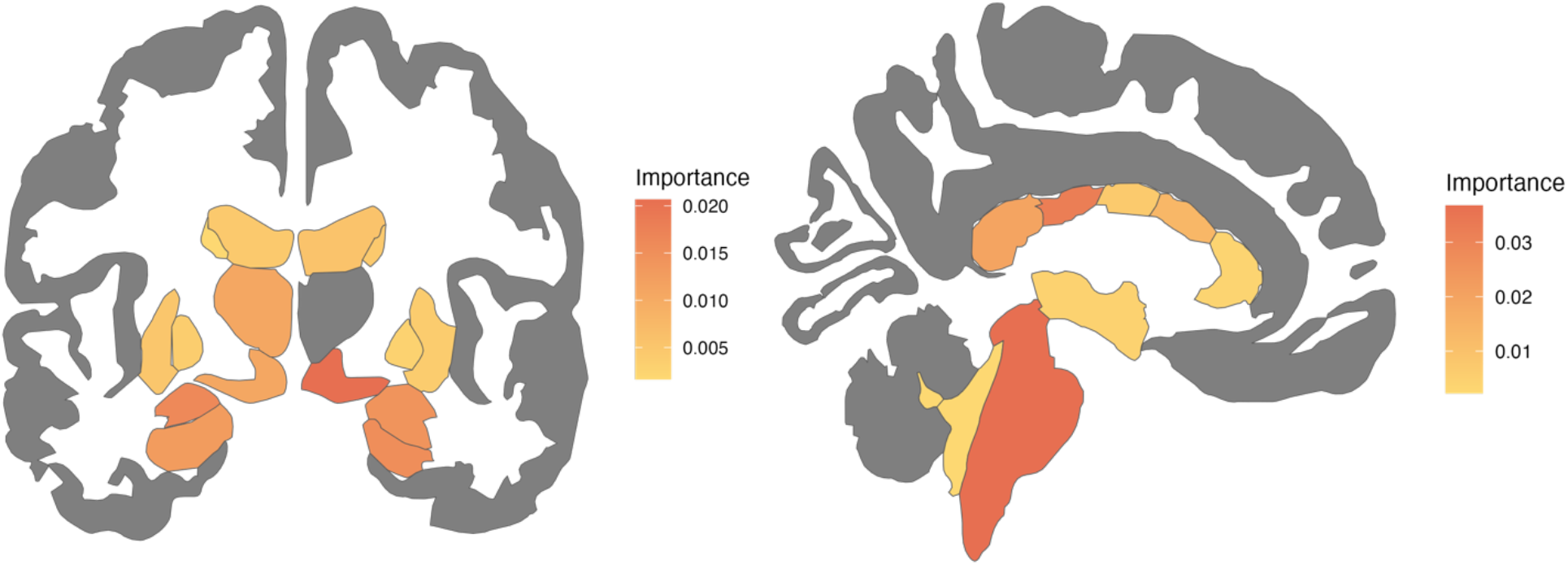
Neuroanatomical contributors to the novel model (baseline). A) Top 20 features ranked by importance to the model, across domains. B) Variable importance of cortical volume features, visualized on 2-dimensional brain. C) Variable importance of cortical area features, visualized on 2-dimensional brain. D) Variable importance of subcortical features, visualized on 2-dimensional brain. Left: Coronal view. Right: Sagittal view.

To contextualize our model features, we compared our findings to those of past adolescent BrainAGE models, including those of Drobonin et al. (2022) and Kelly et al. (2022). While BrainAGE model features cannot always be directly compared due to different brain parcellations and reporting methods, there were a number of high-contributing features in the novel model that are shared with those found in existing work. Shared with Drobonin et al. (2022), we found that the brainstem, superior parietal, isthmus cingulate, ventral diencephalon, amygdala, hippocampus, and posterior corpus callosum were high contributors to the model.

Additionally, we compared our results with those of the model from Kelly et al. (2022) and found that both models shared volumes of frontal, temporal, and parietal regions, the brainstem as high contributors.

#### Follow-Up

As in the baseline model, top contributors to the model were determined using the vip R package (version 0.4.1). The top contributors to the model included the intercranial volume, as well as the volume of the brainstem, mid-posterior and posterior corpus callosum, right isthmus cingulate, right inferior parietal, and left rostral middle frontal regions. Generally, volume measurements contributed more highly to the model than surface area.

Many of these regions were also top contributors to the baseline model, including the brainstem, intercranial volume, and corpus callosum. New high contributors included the right inferior parietal, rostral middle frontal, and lateral ventricle volumes, as well as the caudal anterior cingulate area. Compared to the Drobonin et al. (2022) model, shared regions included the brainstem, corpus callosum, amygdala, inferior parietal, and isthmus cingulate. Shared regions with the Kelly et al. (2022) included the brainstem, ventricles, as well as frontal, temporal, and parietal volumes.

### BrainAGE and Maturational Metrics

#### Baseline

A higher BrainAGE was related to more advanced youth- and parent-report pubertal development (Table 4). For youth-report pubertal development, (*b_youth_* = 0.31, *se* = 0.09, *p* < .001) (Figure 7a). For parent-report pubertal development, (*b_parent_* = 0.19, *se* = 0.06, *p* <.001) (Figure 7b). In addition, a higher/more positive BrainAGE was related to higher cognition scores (*b_cog_* = 0.002, *se* = 0.003, *p* = .005) (Figure 6c).

**Figure 6.**
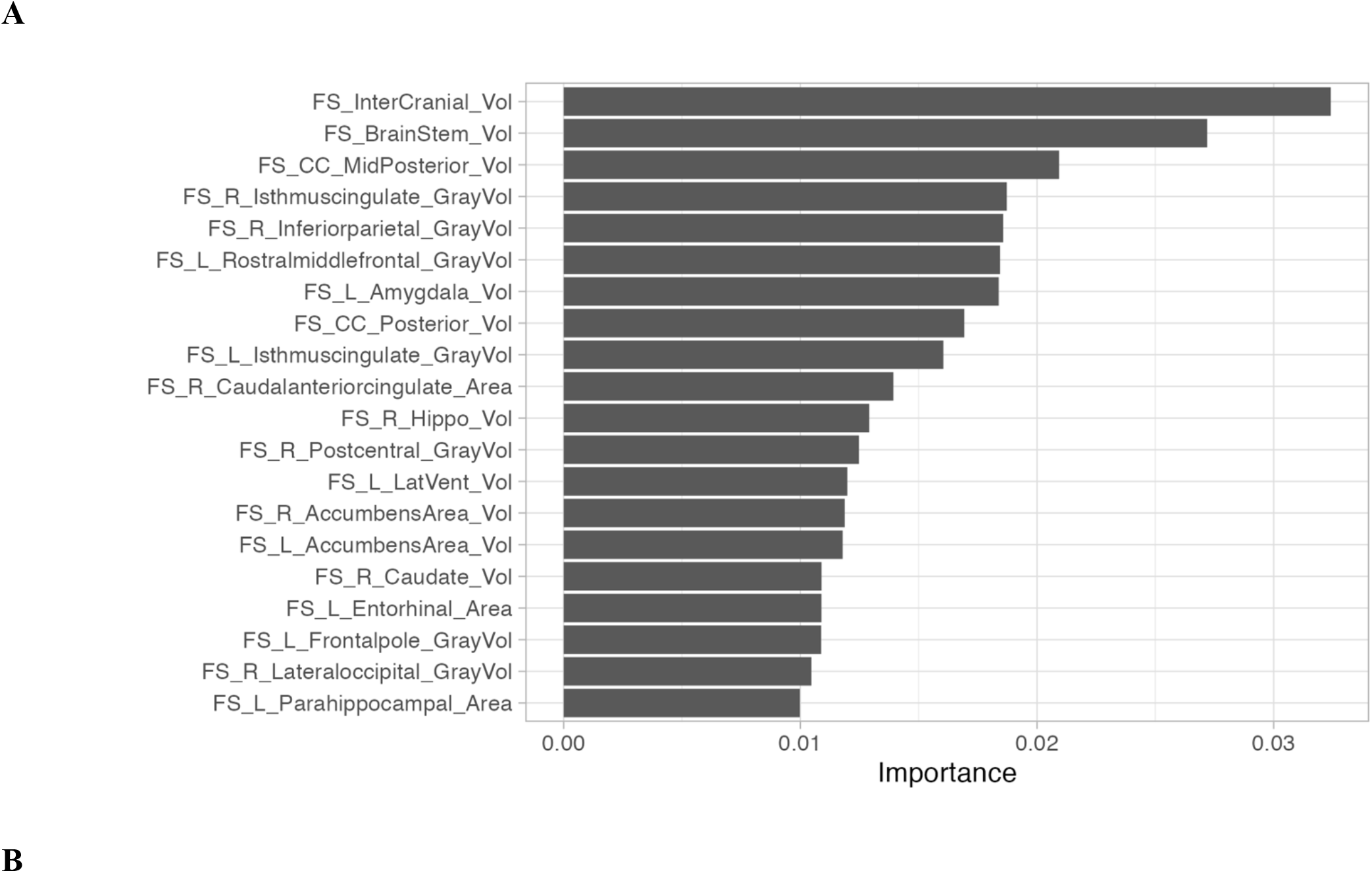
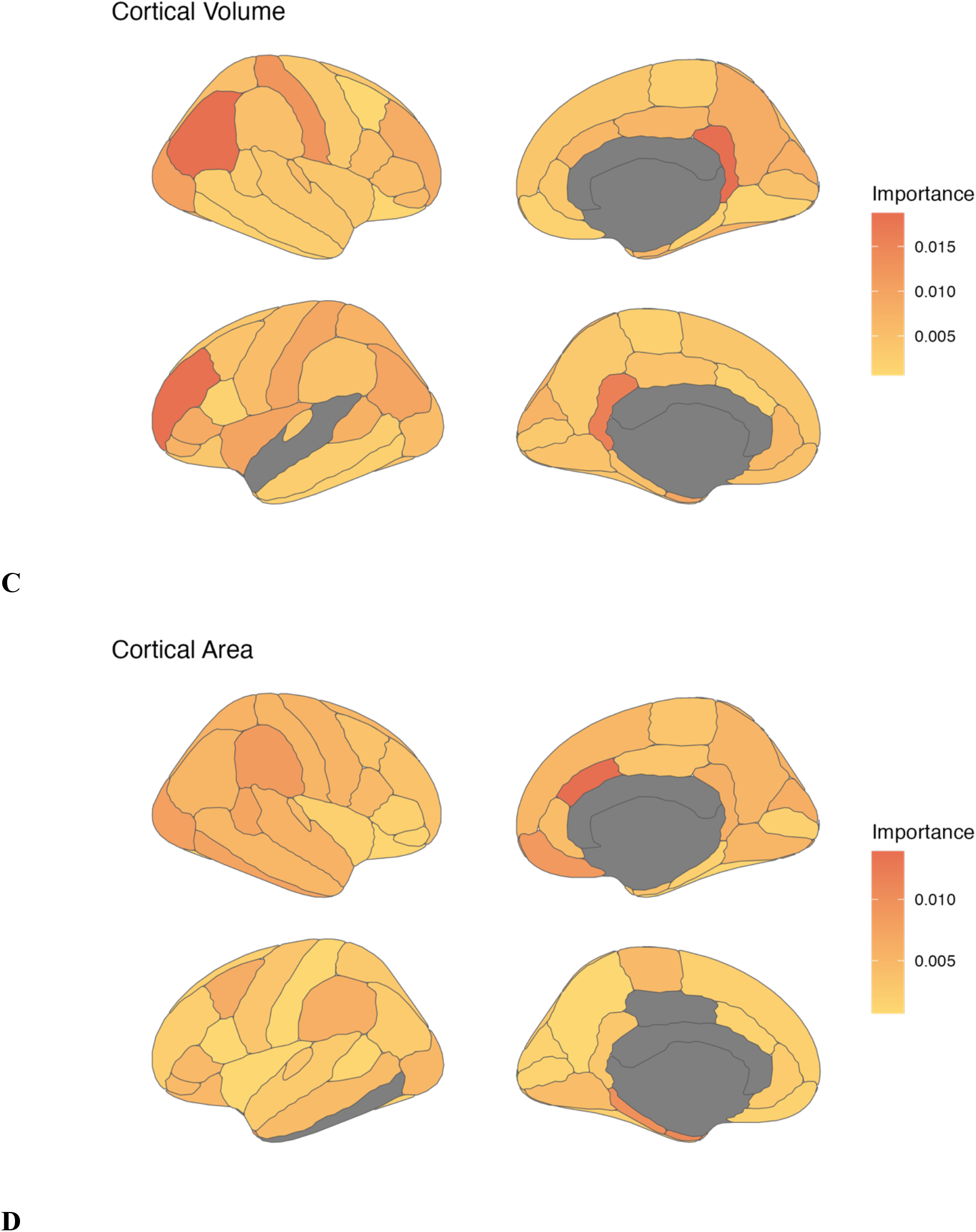
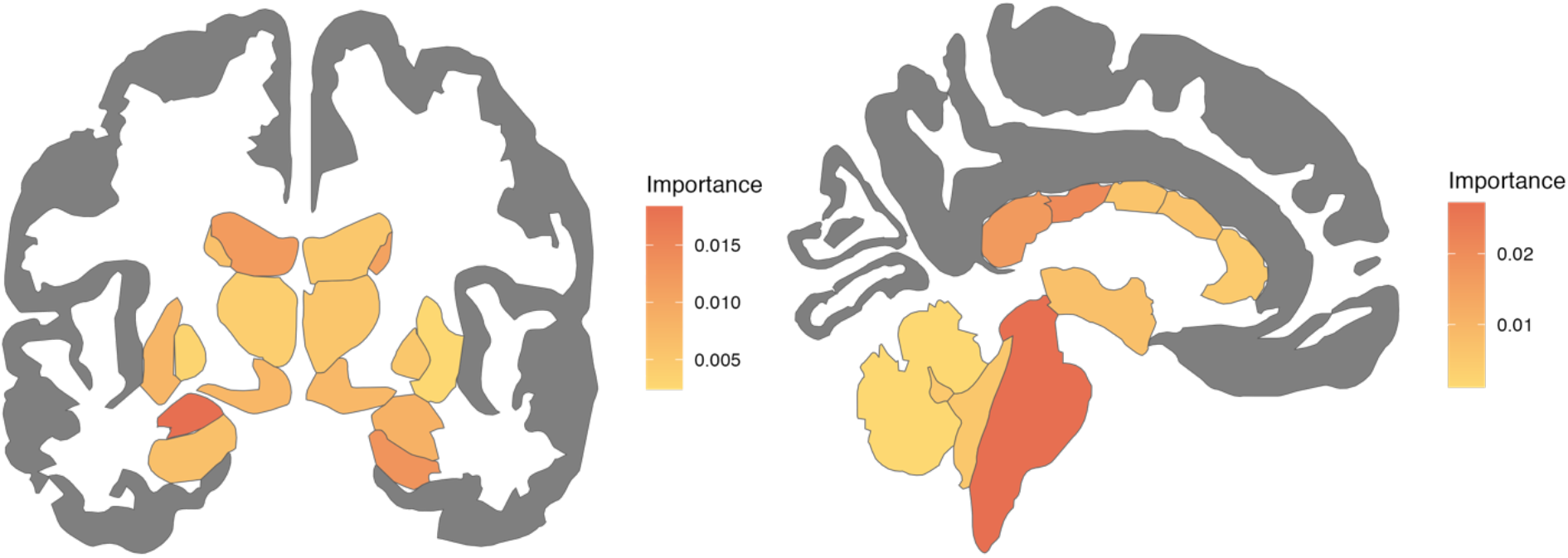
Neuroanatomical contributors to the novel model (follow-up). A) Top 20 features ranked by importance to the model, across domains. B) Variable importance of cortical volume features, visualized on 2-dimensional brain. C) Variable importance of cortical area features, visualized on 2-dimensional brain. D) Variable importance of subcortical features, visualized on 2-dimensional brain. Left: Coronal view. Right: Sagittal view.

**Figure 7.**
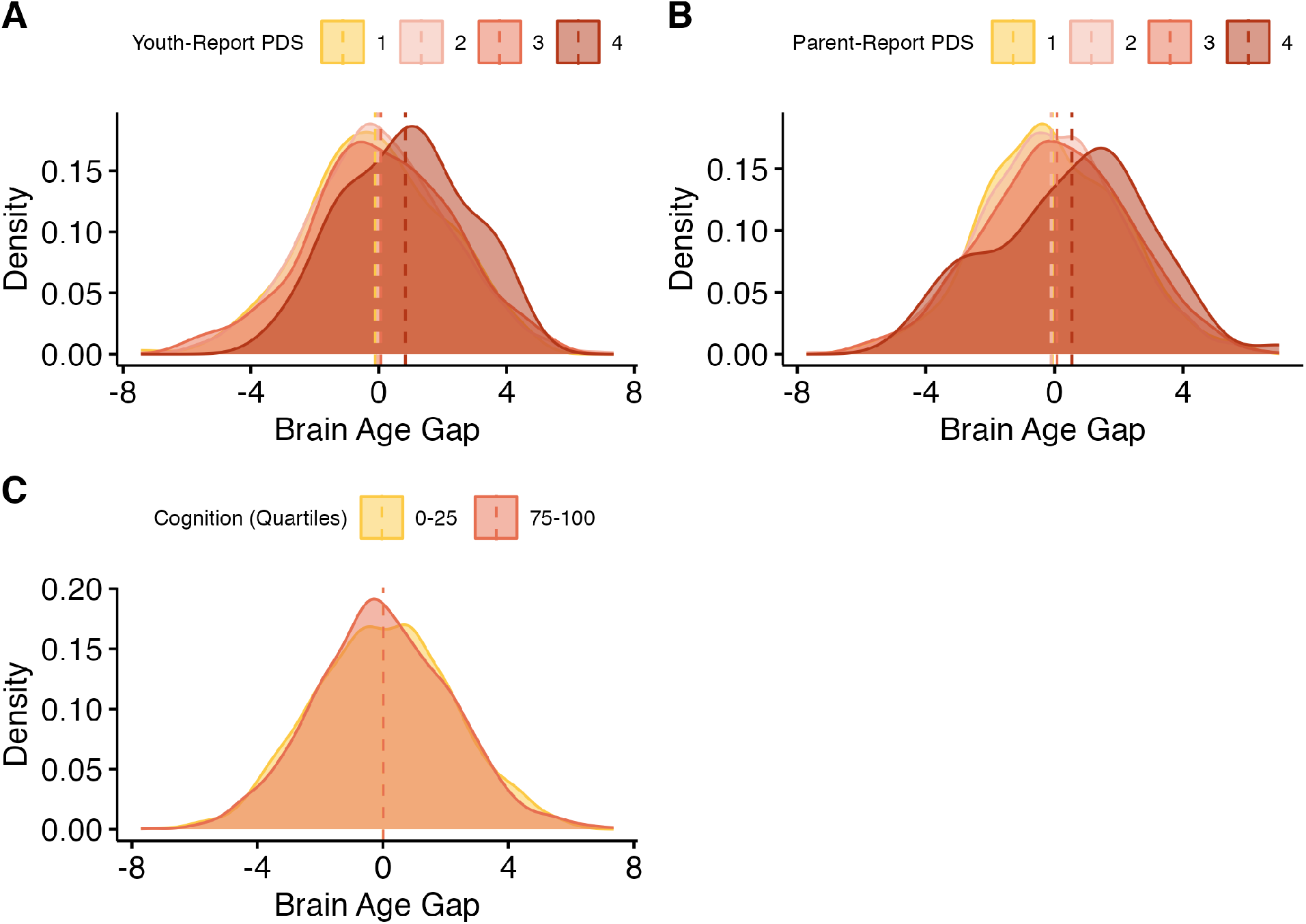
Results - Novel model (baseline). A) Distributions of brain-age gaps, grouped by pubertal stage (youth-report). Means for each group are represented with vertical lines. B) Distributions of brain-age gaps, grouped by pubertal stage (parent-report). Means for each group are represented with vertical lines. C) Distributions of brain-age gaps, grouped by quartile of cognition scores, compared to others in the sample. Only the highest and lowest quartiles are shown. Means for each group are represented with vertical lines.

#### Follow-Up

A higher BrainAGE was related to more advanced youth- and parent-report pubertal development (Table 4). For youth-report pubertal development, (*b_youth_* = 0.39, *se* = 0.08, *p*<.001) (Figure 8a). For parent-report pubertal development, (*b_parent_* = 0.42, *se* = 0.07, *p* <.001) (Figure 8b).

**Figure 8.**
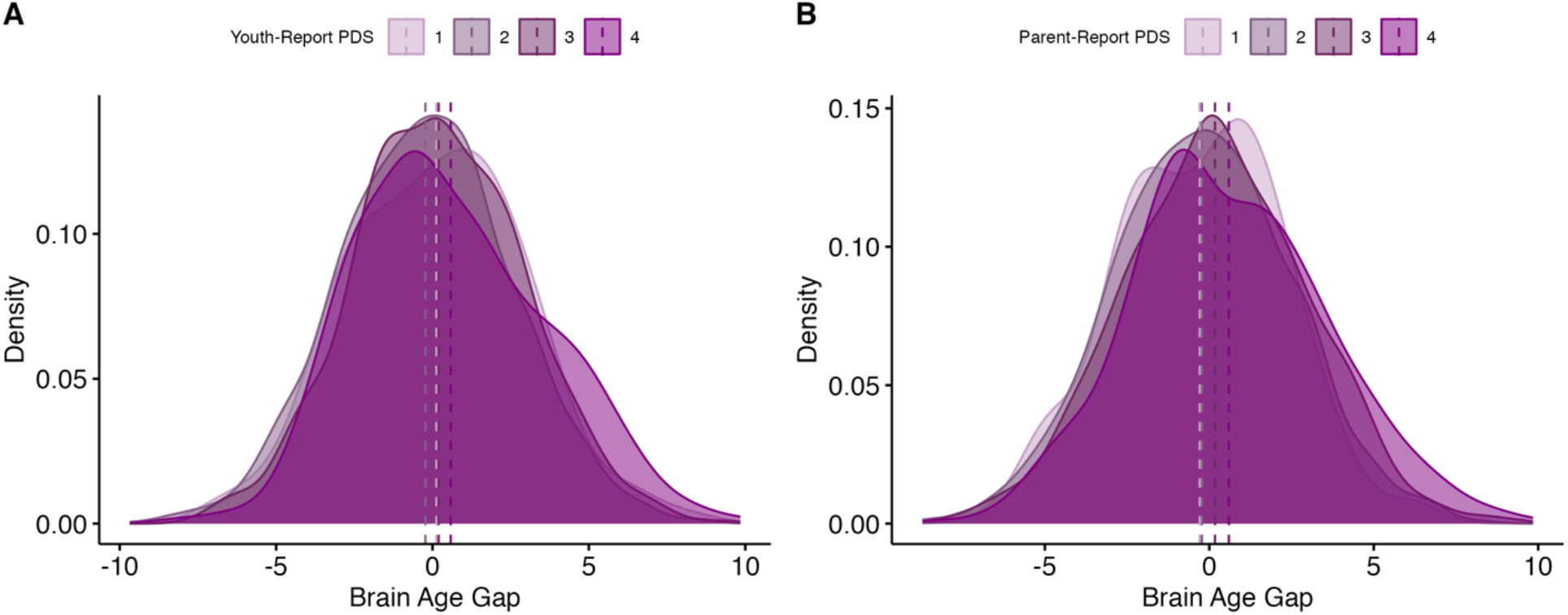
Results - Novel model (follow-up). A) Distributions of brain-age gaps, grouped by pubertal stage (youth-report). Means for each group are represented with vertical lines. B) Distributions of brain-age gaps, grouped by pubertal stage (parent-report). Means for each group are represented with vertical lines.

### Longitudinal Analyses

#### Methods

To examine the stability of BrainAGE measurements over time, we calculated the intraclass correlation (ICC) between BrainAGE at baseline and two-year follow-up for both studies. The ICC quantifies the variance between people and the variance over time, where an ICC of 0 would indicate that all variance is within-person, and an ICC of 1 would indicate that all variance in between-persons (Little et al., 2015). Only participants with BrainAGE estimates at both timepoints were included in these analyses. Using the Study 1 gap predictions, 7,431 participants had data at both timepoints. Using the Study 2 gap predictions, 3,692 participants had estimates at both timepoints.

#### Results

BrainAGE showed moderate to large ICCs across two timepoints in early adolescence. For Study 1, the ICC between BrainAGE estimates was 0.7 (Figure 9a). For Study 2, the ICC between BrainAGE estimates was 0.53 (Figure 9b).

**Figure 9.**
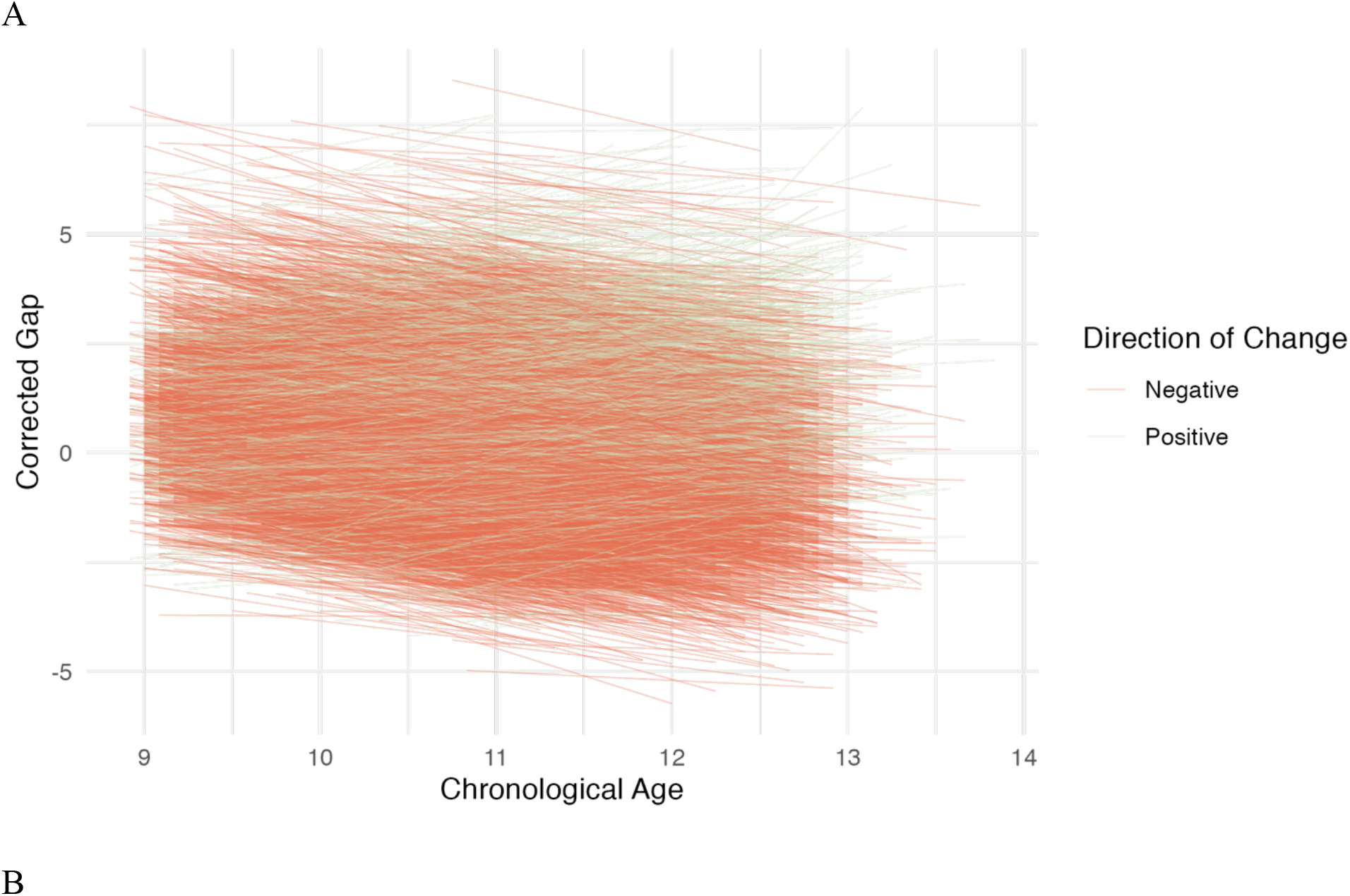
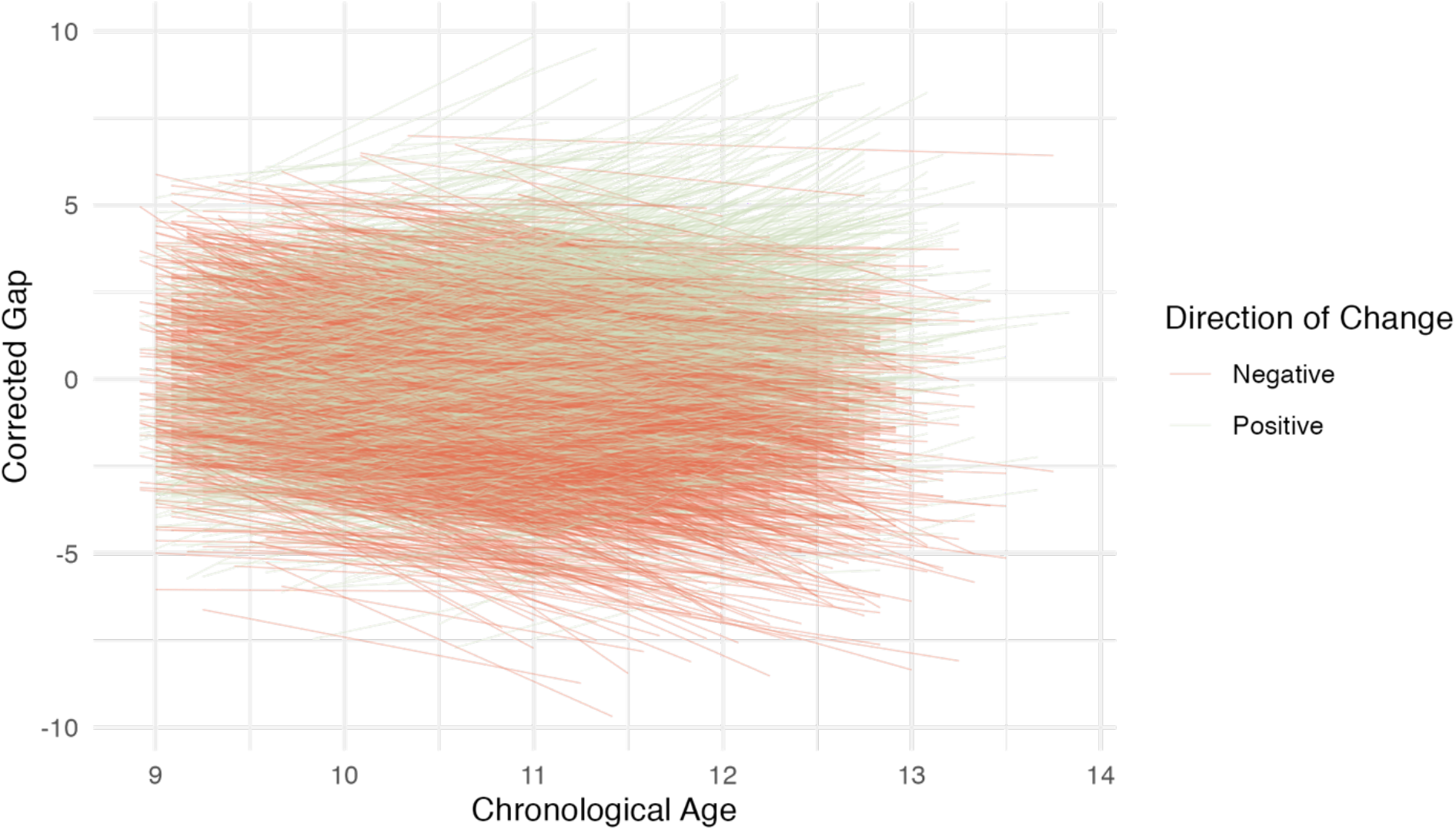
Results – longitudinal changes in gap estimates. A) Individual trajectories of brain-age gaps, where both gaps were predicted using the same model (Study 1). Trajectories are color-coded by whether the estimated gap increased or decreased between timepoints. B). Individual trajectories of brain-age gaps, where the gap at each timepoint was predicted from a timepoint-specific model (Study 2). Trajectories are color-coded by whether the estimated gap increased or decreased between timepoints.

## Discussion

The current study used three BrainAGE models—one previously validated on a wide age range across adolescence and two trained on specific, narrow age ranges in early adolescence— to quantify the relationship between BrainAGE and metrics of pubertal and cognitive development. Using the previously validated model, we established that BrainAGE is positively related to both youth- and parent-report pubertal development, and this relationship holds across two age ranges (9-11, 10-13 years). This association was replicated using our novel models created from independent samples of 9-11 and 10-13 year olds.

The current paper builds on our existing understanding of the relationship between puberty/maturational metrics and BrainAGE in adolescence. Notably, the relationship between pubertal development and BrainAGE was replicated using a model trained on the specific age range of our analysis sample, which featured a lower MAE and reduced age bias. Furthermore, these models and their accompanying training code are available to other researchers to enable further investigation of BrainAGE in early adolescence.

While the relationship between BrainAGE and NIH Toolbox Cognition scores was consistently negatively related across subsamples of estimates derived from the validated model, these results were in conflict with the positive relationship between BrainAGE and cognition scores found in the estimates derived from the novel model. NIH Toolbox Cognition summary scores were not calculated for the two-year follow-up data within ABCD Release 4.0, so it was not possible to examine whether these relationships replicate across waves.

Additionally, we examined longitudinal change in BrainAGE using age-specific models, and found moderate to large ICCs across two timepoints in early adolescence. Stability in BrainAGE was higher when using the same model to predict age at multiple timepoints, compared to using separate models for each timepoint. This indicates that timepoint-specific models may capture more changes that occur in specific developmental periods, but may also be more difficult to interpret between models. Additionally, there was individual variation in the direction of change in BrainAGE, and there were differences in the degree of variation of trajectories between models.

The results of the current study provide initial evidence that BrainAGE tracks with some metrics of maturation, including pubertal development. However, the conflicting results between BrainAGE and cognition lead us to question the utility of these models for non-biological processes. The improvement in MAE between the age groups may be due to the fact that the younger age group was on the lower end/extreme of the age range used to train the model, which might have had fewer participants in the training set than in the older group, so the model would be more accurate for the older age group. Potential sources of the improved MAE (mean absolute error) could be the large amount of data, the narrow age range used in both training and testing (a novel feature of the study), and the use of independent samples of data derived from the same study (ABCD) for training, testing, and prediction in Study 2. In line with our findings, Ball et al. (2017) suggested that image-based models predict most accurately during periods when the rate of anatomical change is greatest. The current study provides a novel BrainAGE model trained on a large sample of early adolescents, which can be used with developmental samples such as ABCD. The novel model is available for use and can be found at https://github.com/LucyWhitmore/BrainAGE-Maturation.

The current study used composite cognition scores which collapse across multiple domains. To accurately assess the relationship between BrainAGE and cognition, we may need to look at individual domains that show dramatic change in early adolescence, such as executive function (Tervo-Clemmens et al., 2022, preprint). While the NIH Toolbox Cognition Battery is generally sensitive to change over our age range of interest, specific domains may be more or less strongly related to BrainAGE (Bauer & Zalazo, 2013).

While the NIH Toolbox is intended to be used over a lifespan age range, measures targeted specifically at constructs that change in early adolescence may provide a clearer view of how BrainAGE relates to cognition in this age range. Furthermore, the lack of complete composite cognition scores at the two-year follow-up limits our ability to examine the relationship between BrainAGE and cognition in multiple age bands. In future work, we may be able to see whether the inconsistent relationship persists in older adolescents.

Additionally, a robust relationship was observed between BrainAGE and puberty across subsamples, waves, models, and reporting methods. These results provide some evidence that BrainAGE tracks with known metrics of pubertal maturation. The relationship between BrainAGE and pubertal development strengthened between waves, possibly due to increased representation of later pubertal stages in the older age range. In future work, these analyses can be extended to track the relationship between BrainAGE and puberty within different age bands in adolescence and illustrate when BrainAGE and puberty are most closely tied. Additionally, youth- and parent-report pubertal development are known to vary in their reliability across age, with parent-report being more reliable at younger ages, and youth-report being more reliable at older ages (Terry et al., 2016). In the current study, we have shown that both reporting methods are consistently linked to BrainAGE. Future work may be able to illustrate when, or if, one method of reporting is no longer associated with BrainAGE.

To make matters more complicated, there are reciprocal connections between many of the constructs that have been related to BrainAGE. Here, we showed that BrainAGE is related to pubertal development, and past work has shown BrainAGE is related to psychopathology as well as adversity and, in adults, early life events such as low birth weight (Drobonin et al., 2022; Vidal-Pineiro et al., 2021). However, earlier pubertal timing has also been related to increased risk for psychopathology, and both psychopathology and early pubertal timing are related to environmental factors (Barendse et al., 2022; Graber et al., 1995; 2013; Mclaughlin et al., 2012). Therefore, it remains difficult to untangle these relationships and mechanisms.

Additionally, more longitudinal BrainAGE work is needed to fully characterize the relationship between BrainAGE and maturational metrics. The current study provides evidence that BrainAGE is positively correlated with pubertal development in early adolescence, but future work should investigate the potentially changing relationship between puberty and BrainAGE in a longitudinal framework, and over a wider range of adolescence.

Our longitudinal analyses showed moderate to large ICCs between BrainAGE at two timepoints, and a larger ICC for BrainAGE predicted by the same model at different waves. However, future work should examine how changes in BrainAGE over time, including changes in direction, relate to outcomes. Furthermore, there is a need for more work considering the impact of age correction and age bias on adolescent BrainAGE models. As discussed earlier, these models are subject to regression to the mean, where younger participants are predicted to be older, and older participants predicted to be younger. While most BrainAGE models are bias corrected, this correction is not perfect. This presents potential issues for longitudinal analyses, where trajectories of BrainAGE may be more likely to be negative when using the same model across waves as younger participants in the model have a larger, positive BrainAGE.

Additionally, many of the features used to train BrainAGE models have huge variability and individual differences in adolescence, and brain maturity is more related to trajectories of brain development than overall values, which is not accounted for in most BrainAGE models (Mills et al., 2021). In current models, a participant may be considered younger or older because of the starting size of their brain, while they are in fact still following a normative developmental trajectory.

In summary, BrainAGE is a powerful tool that has promise as a clinical biomarker and as a tool to inform basic research. However, as BrainAGE gains traction within the field of developmental cognitive neuroscience, we must ensure that we are careful about our interpretations of its predictions and associations and that we are confident in what we are actually measuring when we talk about brain maturity. As the results of brain maturity studies have implications for legal and other policy domains, it is vital that researchers provide contextualization for BrainAGE and be clear about what we can and cannot infer about different forms of maturity. BrainAGE is a powerful and promising tool, but we need to ensure that we have a strong theoretical background before widely incorporating it into studies of adolescent development and maturation.

## Conclusions

The present study demonstrates that BrainAGE is positively correlated with pubertal development in a sample of early adolescents, and that this result is replicable across subsamples, models, and multiple age bands. However, the relationship between BrainAGE and cognition in early adolescence remains unclear. Future work should examine these relationships in later adolescence and utilize cognition measures that are optimized to examine changes across adolescence.

## Contributions

Contributed to conception and design: LBW, KLM

Contributed to acquisition of data: ABCD

Contributed to analysis and interpretation of data: LBW, KLM, SJW

Drafted and/or revised the article: LBW, KLM, SJW

Approved the submitted version for publication: LBW, KLM, SJW

## Acknowledgements

Data used in the preparation of this article were obtained from the Adolescent Brain Cognitive Development (ABCD) Study (https://abcdstudy.org), held in the NIMH Data Archive (NDA). This is a multisite, longitudinal study designed to recruit more than 10,000 children aged 9-10 and follow them over 10 years into early adulthood. The ABCD Study is supported by the National Institutes of Health and additional federal partners under award numbers U01DA041048, U01DA050989, U01DA051016, U01DA041022, U01DA051018, U01DA051037, U01DA050987, U01DA041174, U01DA041106, U01DA041117, U01DA041028, U01DA041134, U01DA050988, U01DA051039, U01DA041156, U01DA041025, U01DA041120, U01DA051038, U01DA041148, U01DA041093, U01DA041089, U24DA041123, U24DA041147. A full list of supporters is available at https://abcdstudy.org/federal-partners.html. A listing of participating sites and a complete listing of the study investigators can be found at https://abcdstudy.org/consortium_members/. ABCD consortium investigators designed and implemented the study and/or provided data but did not necessarily participate in analysis or writing of this report. This manuscript reflects the views of the authors and may not reflect the opinions or views of the NIH or ABCD consortium investigators. The ABCD data repository grows and changes over time. The ABCD data used in this report came from https://dx.doi.org/10.15154/1528926.

We thank Dr. Vlad Drobonin for making his scripts and validated BrainAGE model available for use.

## Data Availability Statement

Analysis scripts can be found at: https://github.com/LucyWhitmore/BrainAGE-Maturation The publicly available existing model, as well as training and testing scripts, can be found at: https://github.com/GitDro/DevelopmentalBrainAge. Data are part of the ABCD 4.0 Data Release, and are available with an NIH Data Use Certification. The 4.0 release can be found at: http://dx.doi.org/10.15154/1523041

## Author Note

No competing interests exist.

## Supplementary Materials

**Supplementary Table 1.**
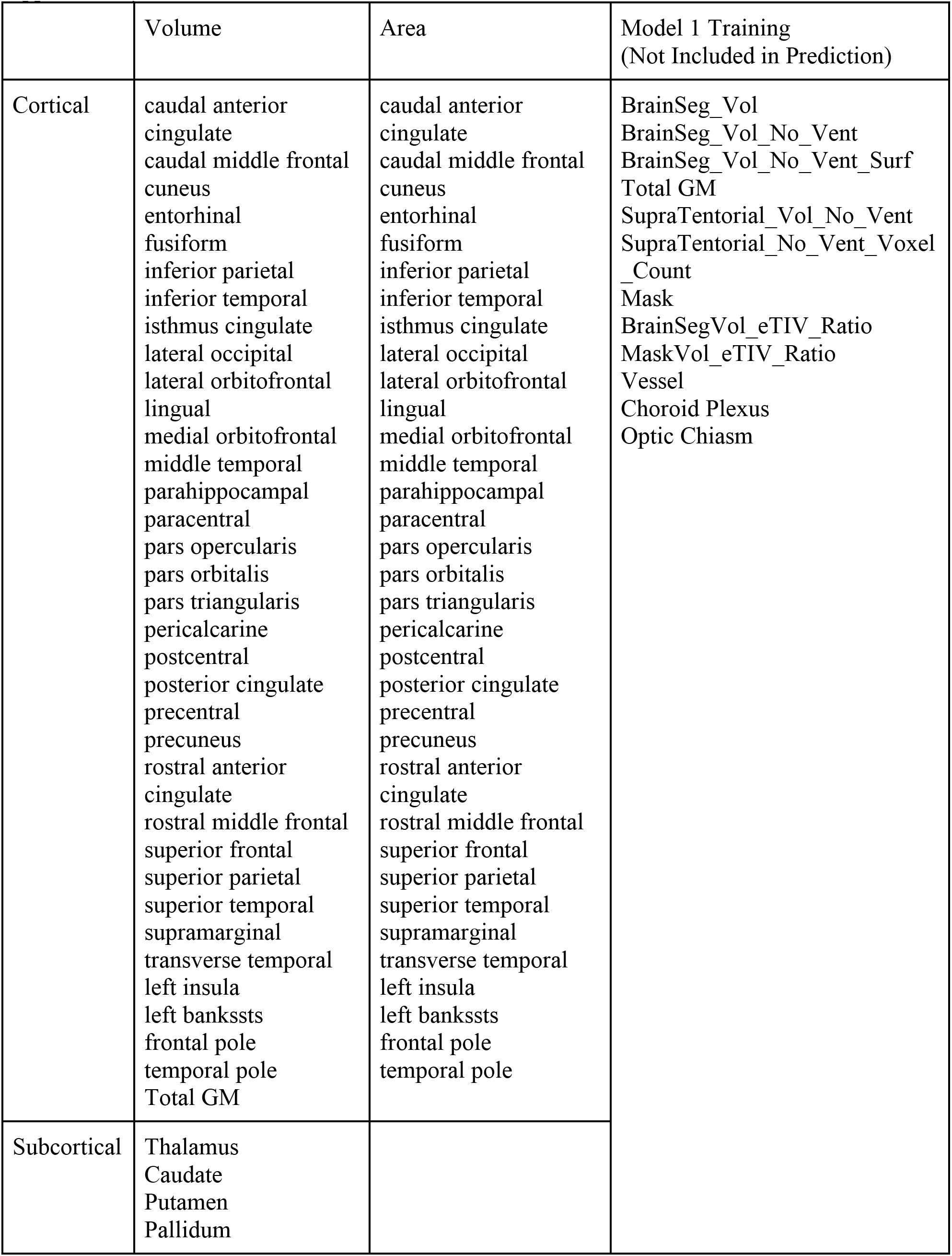
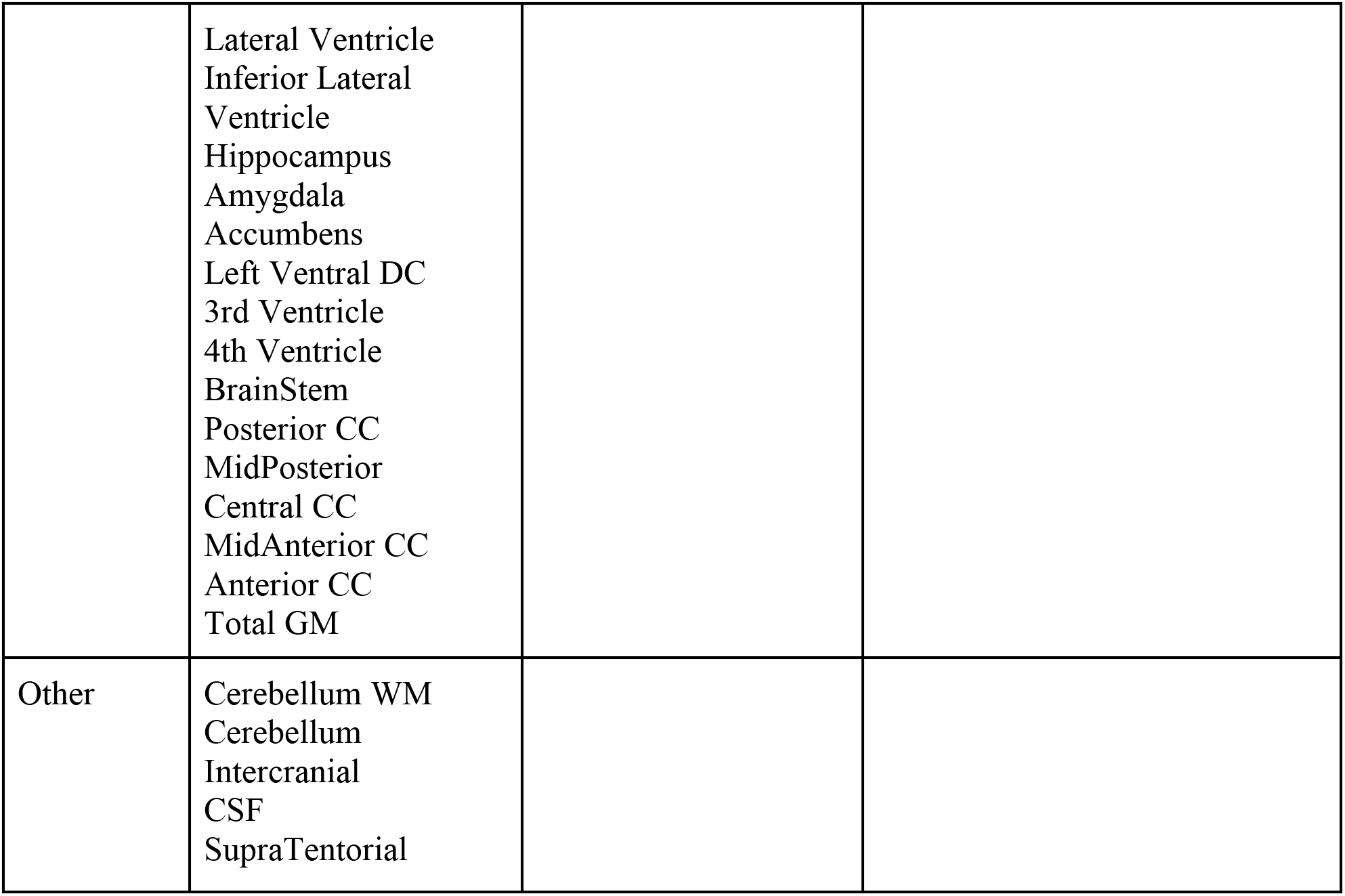
Brain features.

**Supplementary Table 2.**
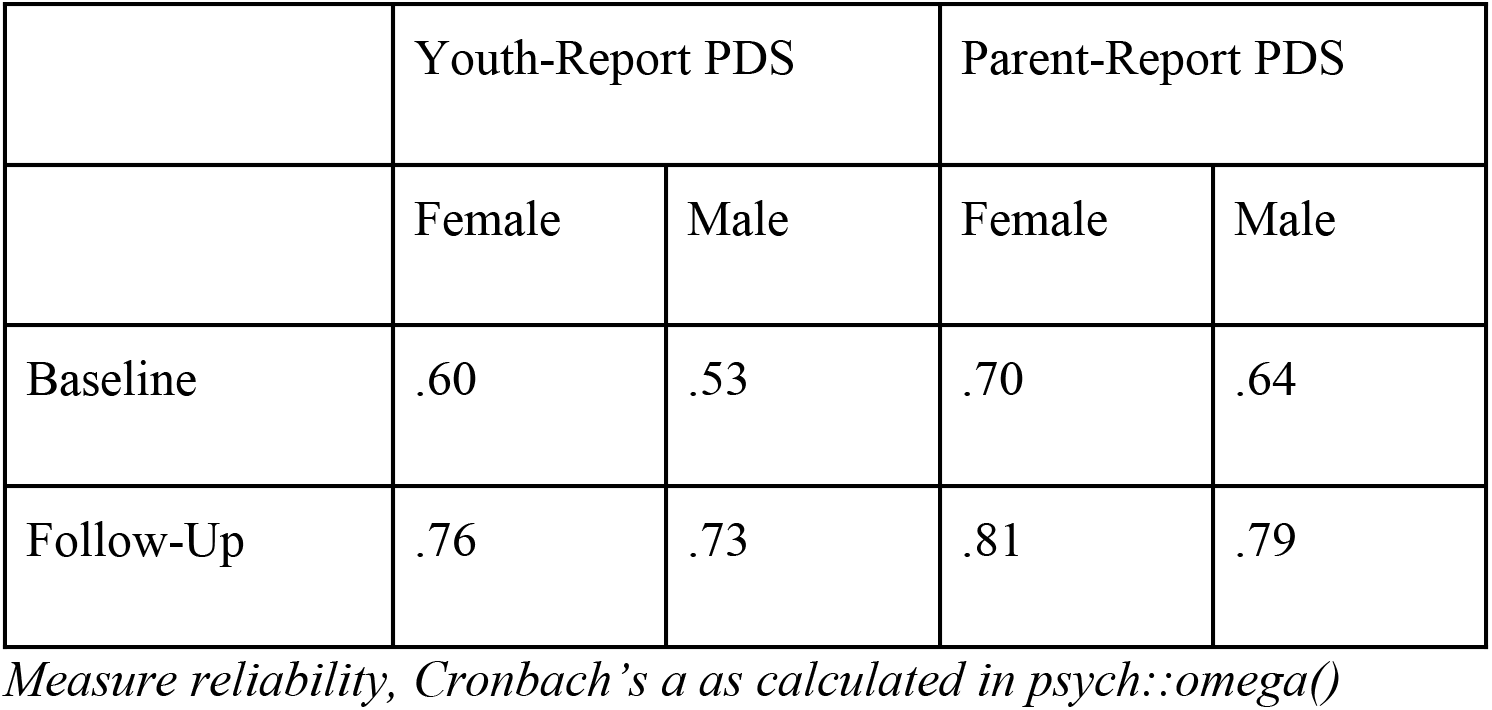
Reliability Measures.

**Supplementary Table 3.**
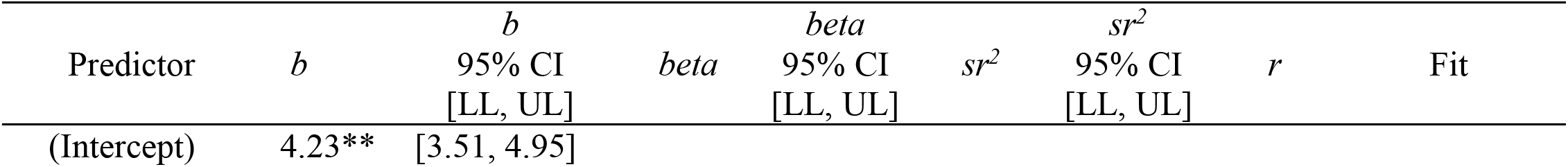
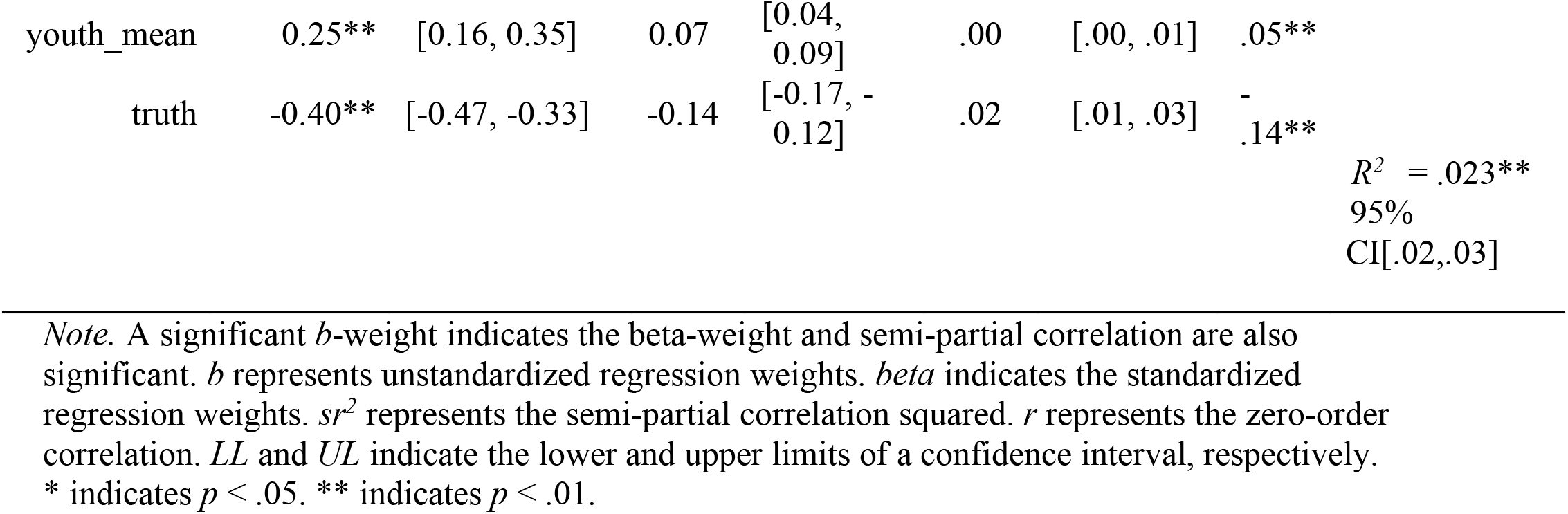
Model 1 – Baseline, Youth-Report PDS. *Regression results using corrected_gap as the criterion*

**Supplementary Table 4.**
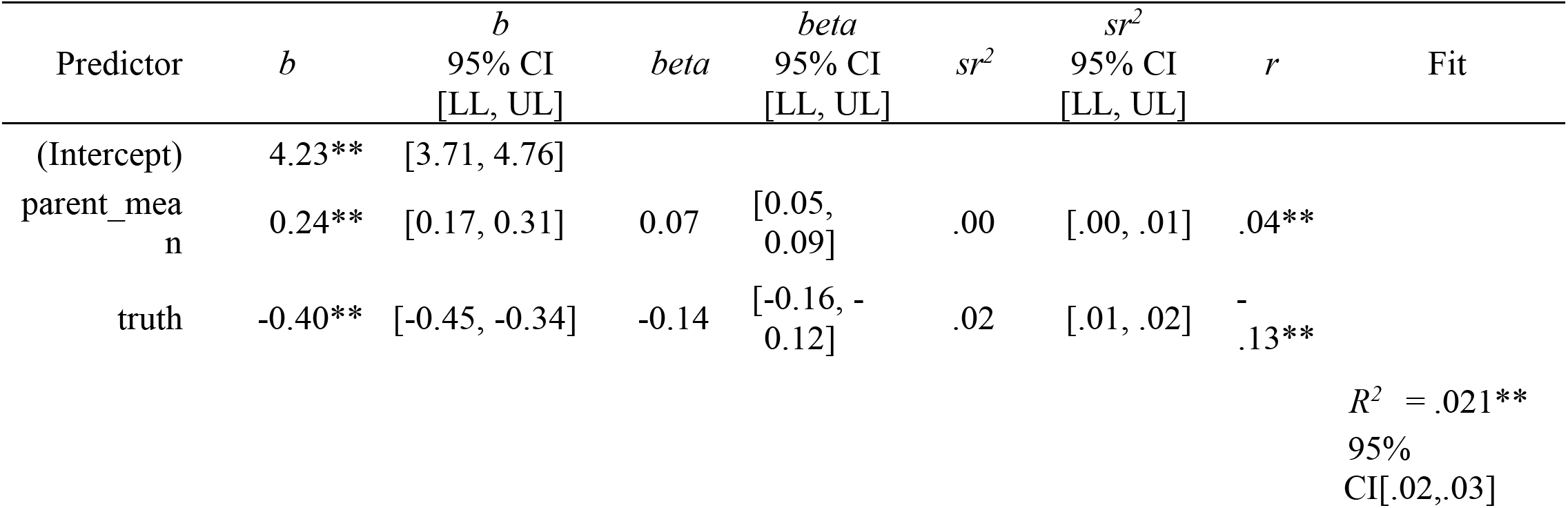
Model 1 – Baseline, Parent-Report PDS. *Regression results using corrected_gap as the criterion*

**Supplementary Table 5.**
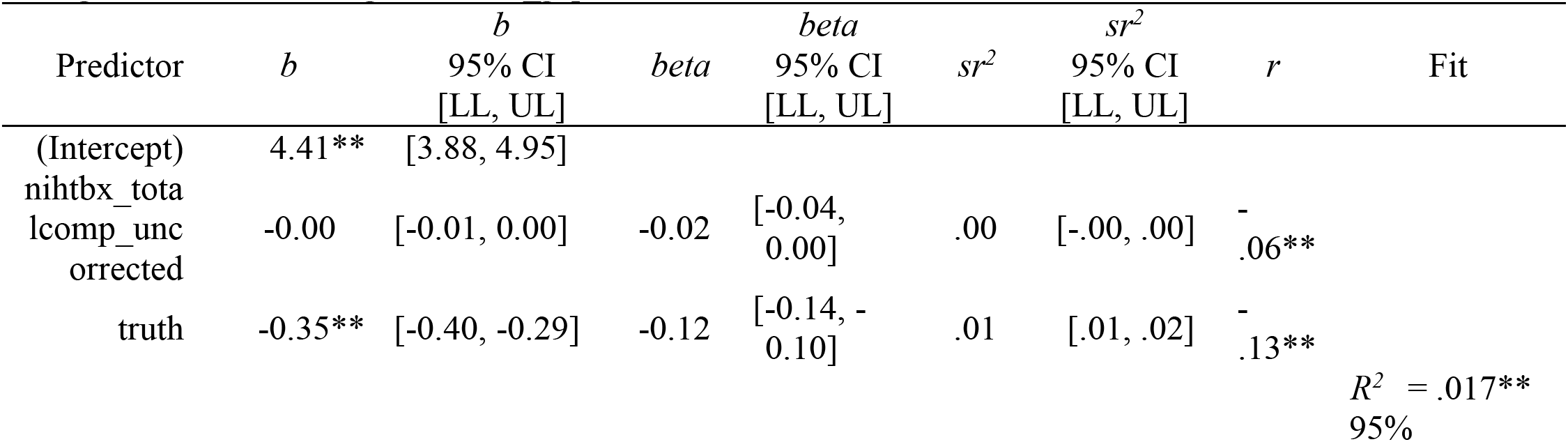
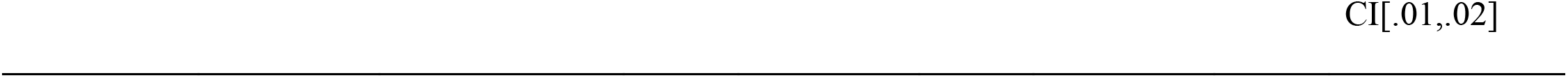
Model 1 – Baseline, Cognition. *Regression results using corrected_gap as the criterion*

**Supplementary Table 6.**
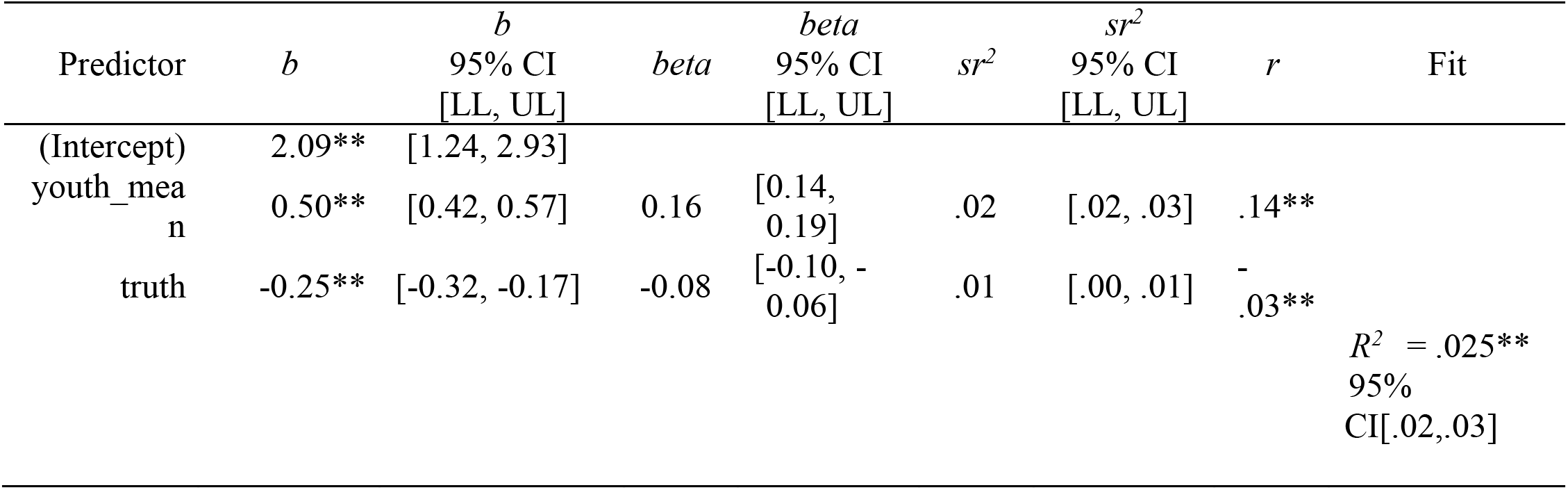
Model 1 – Follow-Up, Youth-Report PDS. *Regression results using corrected_gap as the criterion*

**Supplementary Table 7.**
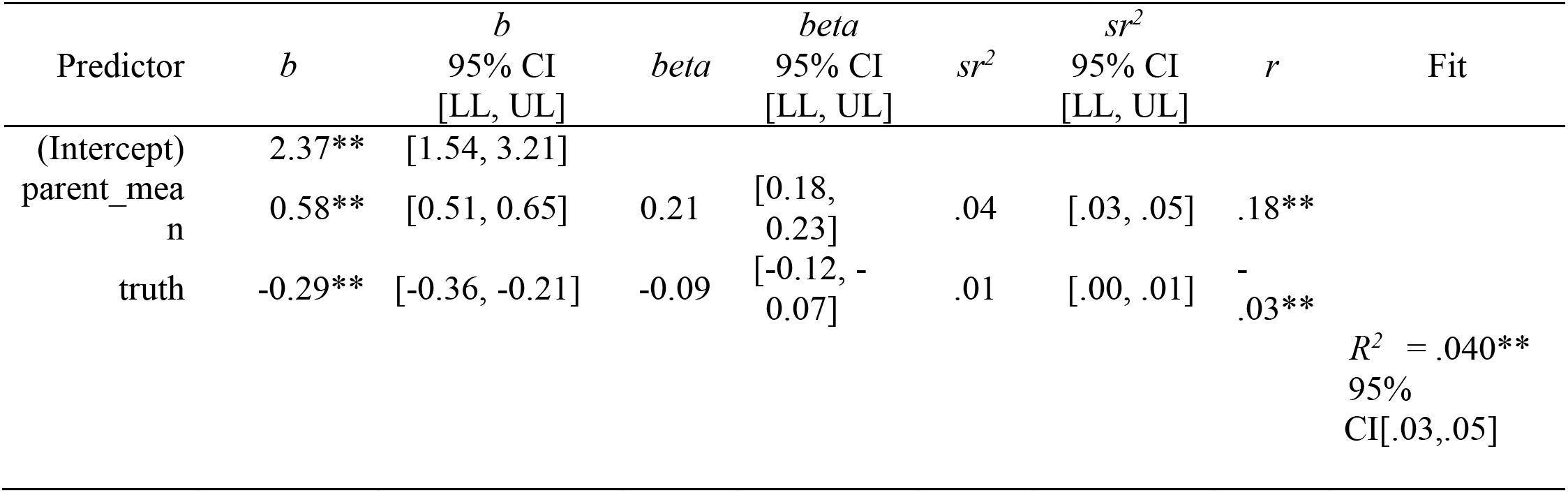
Model 1 – Follow-Up, Parent-Report PDS. *Regression results using corrected_gap as the criterion*

**Supplementary Table 8.**
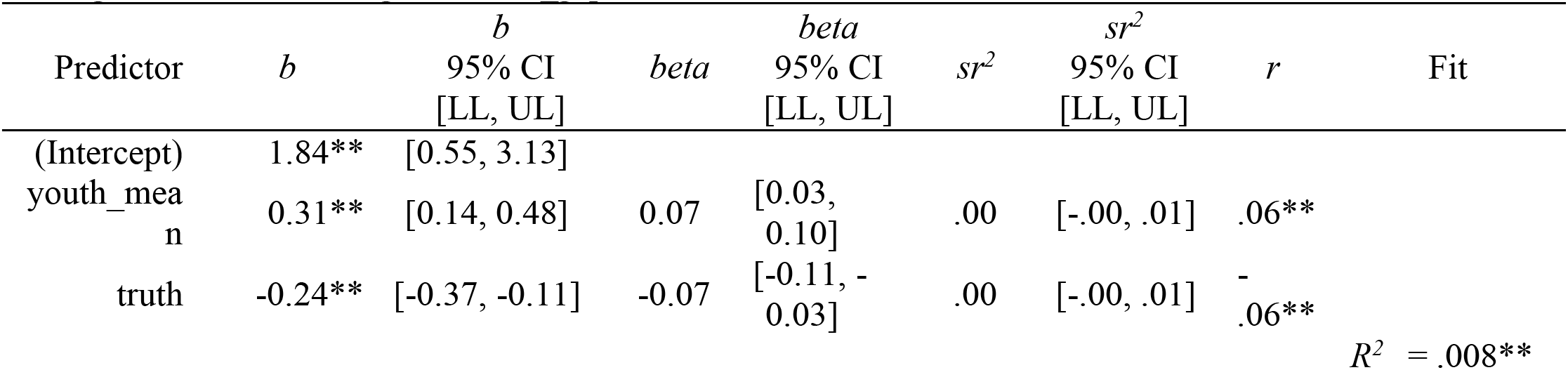
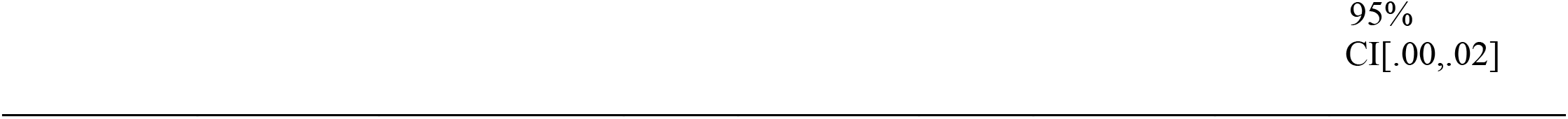
Model 2 – Baseline, Youth-Report PDS. *Regression results using corrected_gap as the criterion*

**Supplementary Table 9.**
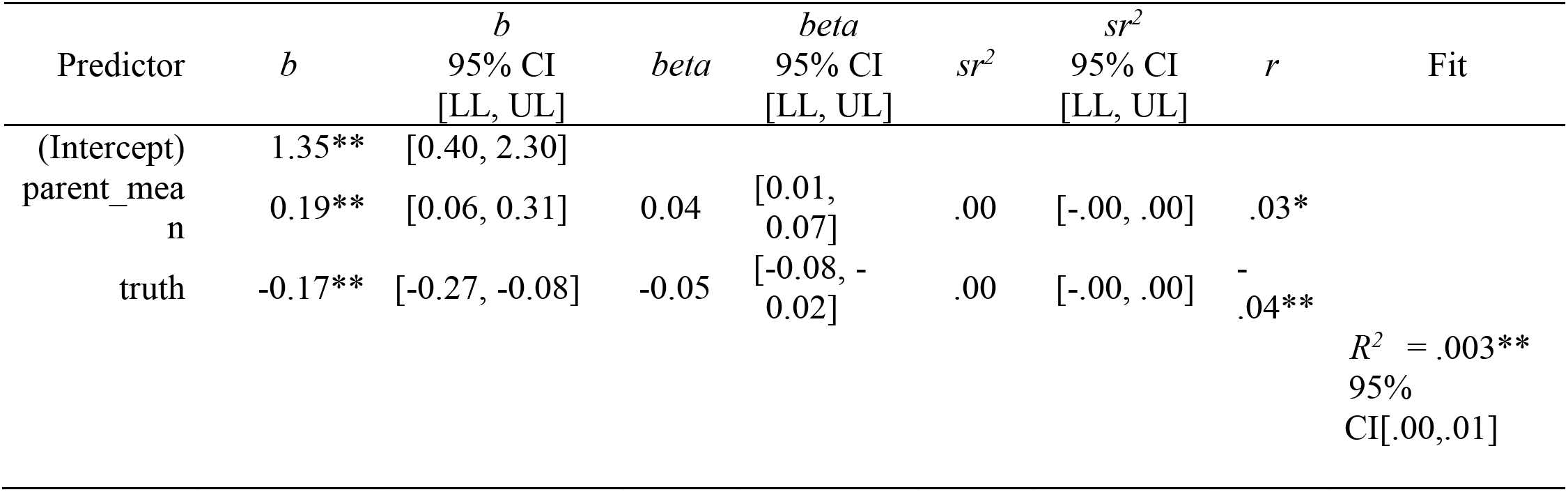
Model 2 – Baseline, Parent-Report PDS. *Regression results using corrected_gap as the criterion*

**Supplementary Table 10.**
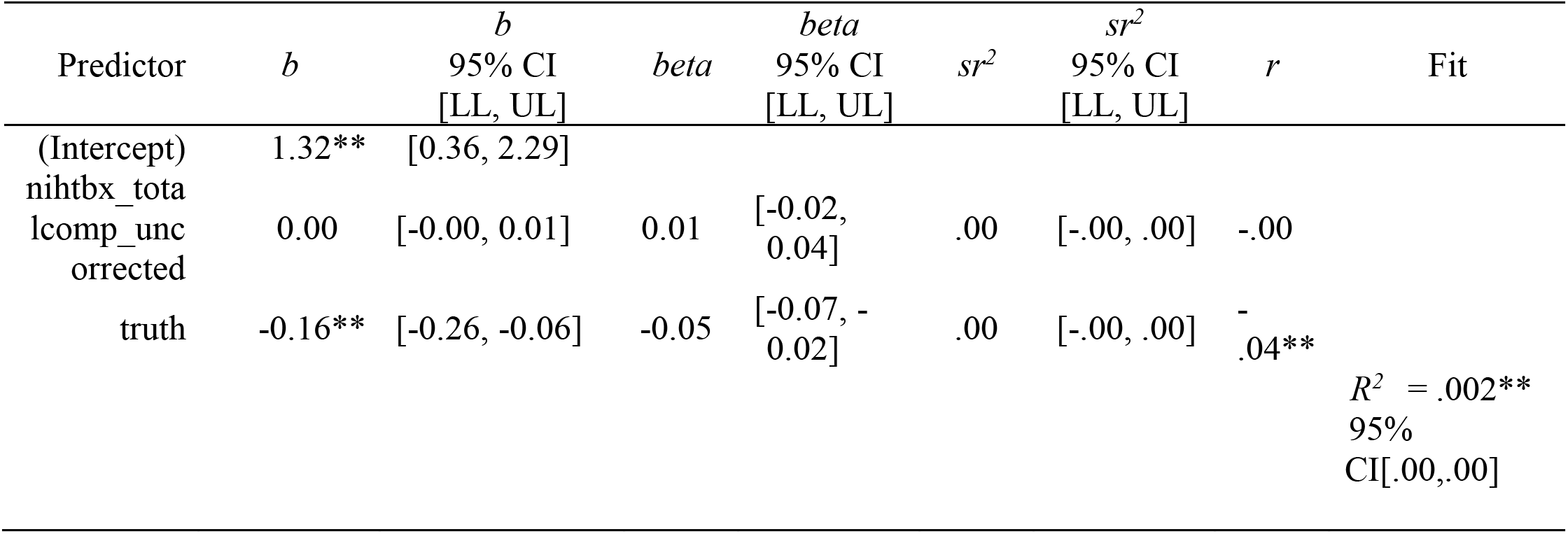
Model 2 – Baseline, Cognition. *Regression results using corrected_gap as the criterion*

**Supplementary Table 11.**
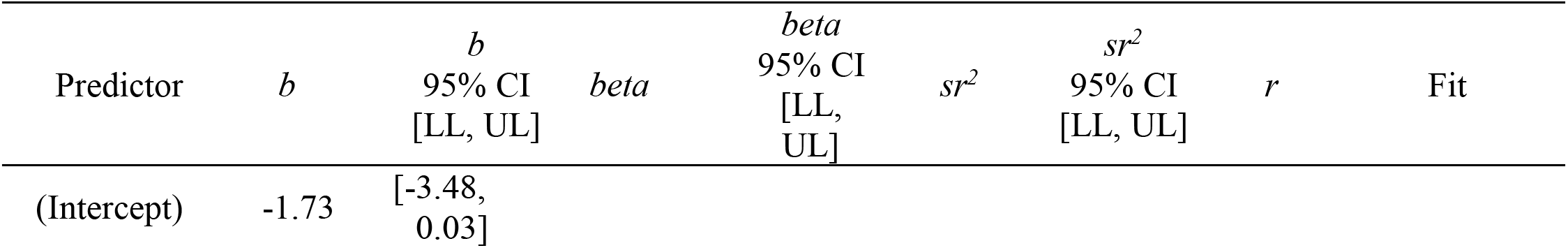
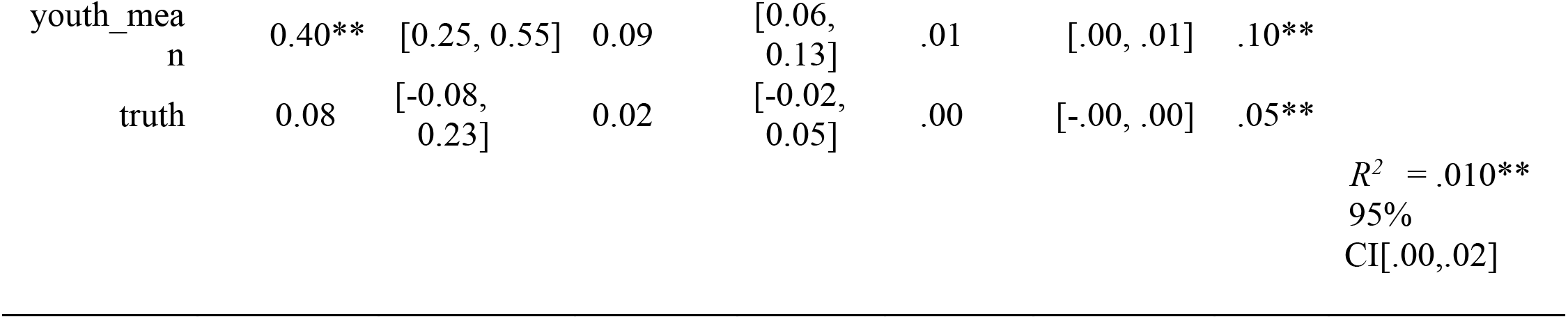
Model 2 – Follow-Up, Youth-Report PDS. *Regression results using corrected_gap as the criterion*

**Supplementary Table 12.**
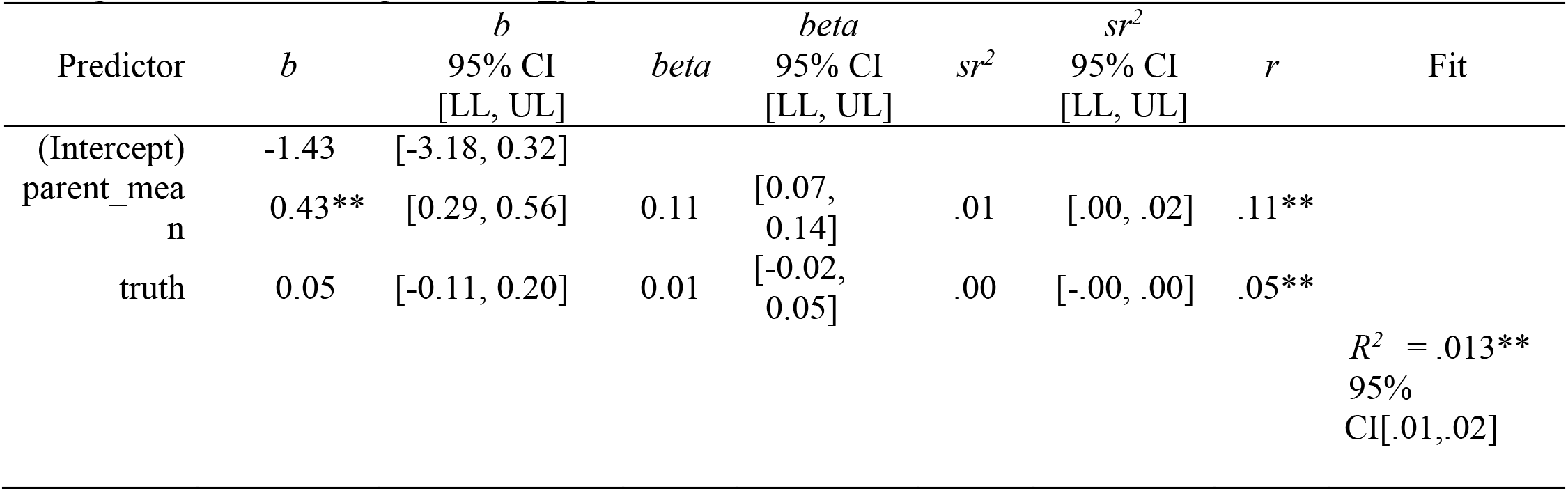
Model 2 – Baseline, Parent-Report PDS. *Regression results using corrected_gap as the criterion*

**Figure S1.**
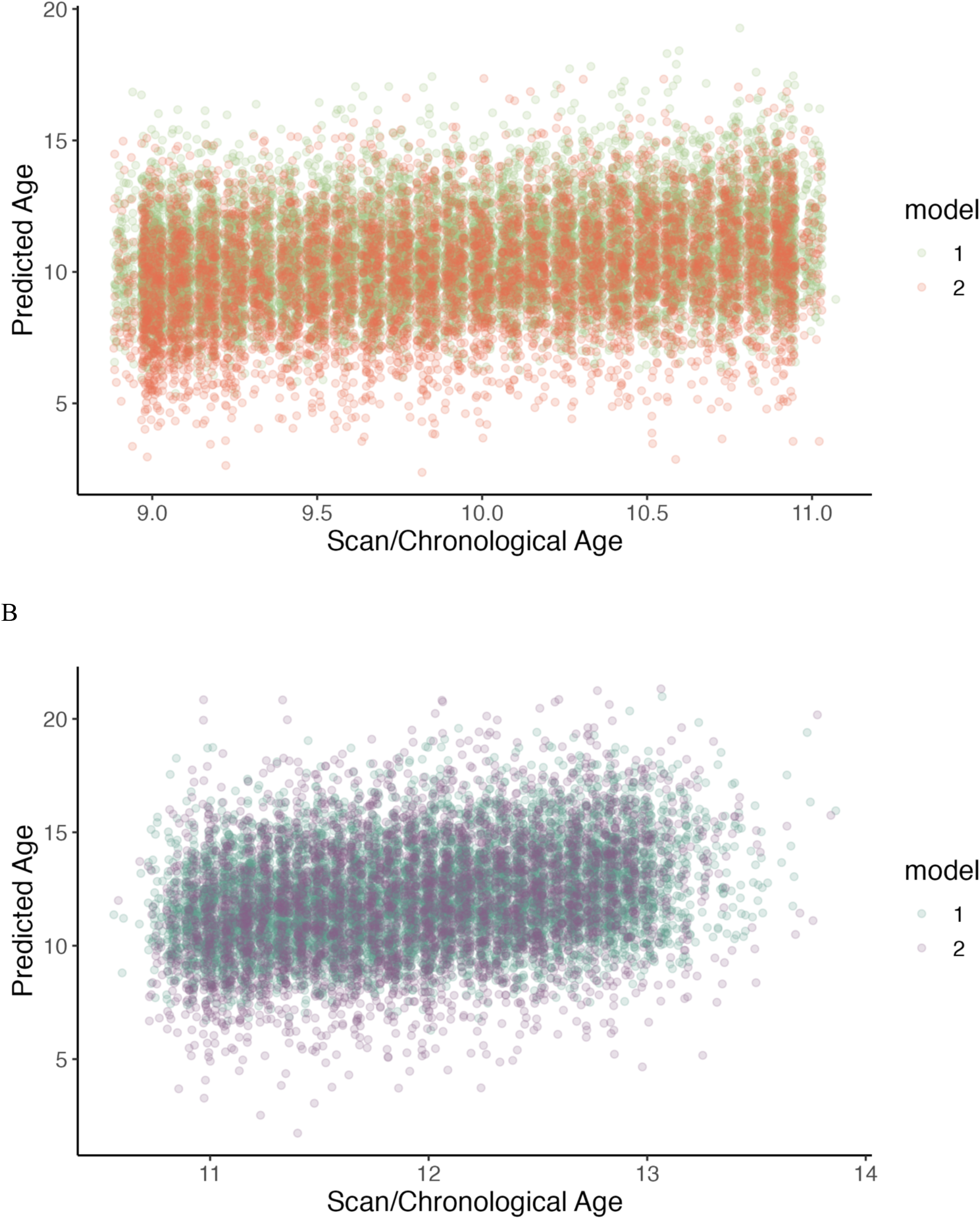
Overlaid model predictions. A) Both baseline models plotted on the same figure. B). Both follow-up models plotted on the same figure.

