## Supplement for "BrainAGE as a measure of maturation during early adolescence"

**Supplementary Materials**

*Supplementary Table 1.* Brain features

| | Volume | Area | Model 1 Training<br>(Not Included in Prediction) |
|---|---|---|---|
| Cortical | caudal anterior cingulate<br>caudal middle frontal<br>cuneus<br>entorhinal<br>fusiform<br>inferior parietal<br>inferior temporal<br>isthmus cingulate<br>lateral occipital<br>lateral orbitofrontal<br>lingual<br>medial orbitofrontal<br>middle temporal<br>parahippocampal<br>paracentral<br>pars opercularis<br>pars orbitalis<br>pars triangularis<br>pericalcarine<br>postcentral<br>posterior cingulate<br>precentral<br>precuneus<br>rostral anterior cingulate<br>rostral middle frontal<br>superior frontal<br>superior parietal<br>superior temporal<br>supramarginal<br>transverse temporal<br>left insula<br>left bankssts<br>frontal pole<br>temporal pole<br>Total GM | caudal anterior cingulate<br>caudal middle frontal<br>cuneus<br>entorhinal<br>fusiform<br>inferior parietal<br>inferior temporal<br>isthmus cingulate<br>lateral occipital<br>lateral orbitofrontal<br>lingual<br>medial orbitofrontal<br>middle temporal<br>parahippocampal<br>paracentral<br>pars opercularis<br>pars orbitalis<br>pars triangularis<br>pericalcarine<br>postcentral<br>posterior cingulate<br>precentral<br>precuneus<br>rostral anterior cingulate<br>rostral middle frontal<br>superior frontal<br>superior parietal<br>superior temporal<br>supramarginal<br>transverse temporal<br>left insula<br>left bankssts<br>frontal pole<br>temporal pole | BrainSeg_Vol<br>BrainSeg_Vol_No_Vent<br>BrainSeg_Vol_No_Vent_Surf<br>Total GM<br>SupraTentorial_Vol_No_Vent<br>SupraTentorial_No_Vent_Voxel_Count<br>Mask<br>BrainSegVol_eTIV_Ratio<br>MaskVol_eTIV_Ratio<br>Vessel<br>Choroid Plexus<br>Optic Chiasm |

| | |
|---|---|
| Subcortical | Thalamus<br>Caudate<br>Putamen<br>Pallidum<br>Lateral Ventricle<br>Inferior Lateral Ventricle<br>Hippocampus<br>Amygdala<br>Accumbens<br>Left Ventral DC<br>3rd Ventricle<br>4th Ventricle<br>BrainStem<br>Posterior CC<br>MidPosterior<br>Central CC<br>MidAnterior CC<br>Anterior CC<br>Total GM |
| Other | Cerebellum WM<br>Cerebellum<br>Intercranial<br>CSF<br>SupraTentorial |

Supplementary Table 2. Reliability Measures

| | Youth-Report PDS | | Parent-Report PDS | |
|---|---|---|---|---|
| | Female | Male | Female | Male |
| Baseline | .60 | .53 | .70 | .64 |
| Follow-Up | .76 | .73 | .81 | .79 |

*Measure reliability, Cronbach's a as calculated in psych::omega()*

Supplementary Table 3. Model 1 – Baseline, Youth-Report PDS
*Regression results using corrected_gap as the criterion*

| Predictor | b | b 95% CI [LL, UL] | beta | beta 95% CI [LL, UL] | $sr^2$ | $sr^2$ 95% CI [LL, UL] | r | Fit |
|---|---|---|---|---|---|---|---|---|
| (Intercept) | 4.23** | [3.51, 4.95] | | | | | | |
| youth_mean | 0.25** | [0.16, 0.35] | 0.07 | [0.04, 0.09] | .00 | [.00, .01] | .05** | |
| truth | -0.40** | [-0.47, -0.33] | -0.14 | [-0.17, -0.12] | .02 | [.01, .03] | -.14** | |
| | | | | | | | | $R^2$ = .023** 95% CI[.02,.03] |

*Note.* A significant *b*-weight indicates the beta-weight and semi-partial correlation are also significant. *b* represents unstandardized regression weights. *beta* indicates the standardized regression weights. $sr^2$ represents the semi-partial correlation squared. *r* represents the zero-order correlation. *LL* and *UL* indicate the lower and upper limits of a confidence interval, respectively. * indicates $p < .05$. ** indicates $p < .01$.

Supplementary Table 4. Model 1 – Baseline, Parent-Report PDS
*Regression results using corrected_gap as the criterion*

| Predictor | b | b 95% CI [LL, UL] | beta | beta 95% CI [LL, UL] | $sr^2$ | $sr^2$ 95% CI [LL, UL] | r | Fit |
|---|---|---|---|---|---|---|---|---|
| (Intercept) | 4.23** | [3.71, 4.76] | | | | | | |
| parent_mean | 0.24** | [0.17, 0.31] | 0.07 | [0.05, 0.09] | .00 | [.00, .01] | .04** | |
| truth | -0.40** | [-0.45, -0.34] | -0.14 | [-0.16, -0.12] | .02 | [.01, .02] | -.13** | |
| | | | | | | | | $R^2$ = .021** 95% CI[.02,.03] |

Supplementary Table 5. Model 1 – Baseline, Cognition
*Regression results using corrected_gap as the criterion*

| Predictor | b | b 95% CI [LL, UL] | beta | beta 95% CI [LL, UL] | $sr^2$ | $sr^2$ 95% CI [LL, UL] | r | Fit |
|---|---|---|---|---|---|---|---|---|
| (Intercept) | 4.41** | [3.88, 4.95] | | | | | | |
| nihtbx_totalcomp_uncorrected | -0.00 | [-0.01, 0.00] | -0.02 | [-0.04, 0.00] | .00 | [-.00, .00] | -.06** | |

| | | | | | | | | |
|---|---|---|---|---|---|---|---|---|
| truth | -0.35** | [-0.40, -0.29] | -0.12 | [-0.14, -0.10] | .01 | [.01, .02] | -.13** | |
| | | | | | | | | $R^2$ = .017** 95% CI[.01,.02] |

Supplementary Table 6. Model 1 – Follow-Up, Youth-Report PDS
*Regression results using corrected_gap as the criterion*

| Predictor | *b* | *b* 95% CI [LL, UL] | *beta* | *beta* 95% CI [LL, UL] | $sr^2$ | $sr^2$ 95% CI [LL, UL] | *r* | Fit |
|---|---|---|---|---|---|---|---|---|
| (Intercept) | 2.09** | [1.24, 2.93] | | | | | | |
| youth_mean | 0.50** | [0.42, 0.57] | 0.16 | [0.14, 0.19] | .02 | [.02, .03] | .14** | |
| truth | -0.25** | [-0.32, -0.17] | -0.08 | [-0.10, -0.06] | .01 | [.00, .01] | -.03** | |
| | | | | | | | | $R^2$ = .025** 95% CI[.02,.03] |

Supplementary Table 7. Model 1 – Follow-Up, Parent-Report PDS
*Regression results using corrected_gap as the criterion*

| Predictor | *b* | *b* 95% CI [LL, UL] | *beta* | *beta* 95% CI [LL, UL] | $sr^2$ | $sr^2$ 95% CI [LL, UL] | *r* | Fit |
|---|---|---|---|---|---|---|---|---|
| (Intercept) | 2.37** | [1.54, 3.21] | | | | | | |
| parent_mean | 0.58** | [0.51, 0.65] | 0.21 | [0.18, 0.23] | .04 | [.03, .05] | .18** | |
| truth | -0.29** | [-0.36, -0.21] | -0.09 | [-0.12, -0.07] | .01 | [.00, .01] | -.03** | |
| | | | | | | | | $R^2$ = .040** 95% CI[.03,.05] |

Supplementary Table 8. Model 2 – Baseline, Youth-Report PDS
*Regression results using corrected_gap as the criterion*

| Predictor | *b* | *b* 95% CI [LL, UL] | *beta* | *beta* 95% CI [LL, UL] | $sr^2$ | $sr^2$ 95% CI [LL, UL] | *r* | Fit |
|---|---|---|---|---|---|---|---|---|
| (Intercept) | 1.84** | [0.55, 3.13] | | | | | | |
| youth_mea | 0.31** | [0.14, 0.48] | 0.07 | [0.03, | .00 | [-.00, .01] | .06** | |

| | | | | | | | | |
|---|---|---|---|---|---|---|---|---|
| n | | | | 0.10] | | | | |
| truth | -0.24** | [-0.37, -0.11] | -0.07 | [-0.11, -0.03] | .00 | [-.00, .01] | -.06** | |
| | | | | | | | | $R^2$ = .008** 95% CI[.00,.02] |

Supplementary Table 9. Model 2 – Baseline, Parent-Report PDS

*Regression results using corrected_gap as the criterion*

| Predictor | *b* | *b* 95% CI [LL, UL] | *beta* | *beta* 95% CI [LL, UL] | $sr^2$ | $sr^2$ 95% CI [LL, UL] | *r* | Fit |
|---|---|---|---|---|---|---|---|---|
| (Intercept) | 1.35** | [0.40, 2.30] | | | | | | |
| parent_mean | 0.19** | [0.06, 0.31] | 0.04 | [0.01, 0.07] | .00 | [-.00, .00] | .03* | |
| truth | -0.17** | [-0.27, -0.08] | -0.05 | [-0.08, -0.02] | .00 | [-.00, .00] | -.04** | |
| | | | | | | | | $R^2$ = .003** 95% CI[.00,.01] |

Supplementary Table 10. Model 2 – Baseline, Cognition

*Regression results using corrected_gap as the criterion*

| Predictor | *b* | *b* 95% CI [LL, UL] | *beta* | *beta* 95% CI [LL, UL] | $sr^2$ | $sr^2$ 95% CI [LL, UL] | *r* | Fit |
|---|---|---|---|---|---|---|---|---|
| (Intercept) | 1.32** | [0.36, 2.29] | | | | | | |
| nihtbx_totalcomp_uncorrected | 0.00 | [-0.00, 0.01] | 0.01 | [-0.02, 0.04] | .00 | [-.00, .00] | -.00 | |
| truth | -0.16** | [-0.26, -0.06] | -0.05 | [-0.07, -0.02] | .00 | [-.00, .00] | -.04** | |
| | | | | | | | | $R^2$ = .002** 95% CI[.00,.00] |

Supplementary Table 11. Model 2 – Follow-Up, Youth-Report PDS

*Regression results using corrected_gap as the criterion*

| Predictor | *b* | *b* 95% CI [LL, UL] | *beta* | *beta* 95% CI [LL, | $sr^2$ | $sr^2$ 95% CI [LL, UL] | *r* | Fit |
|---|---|---|---|---|---|---|---|---|

| | | | | UL] | | | | |
|---|---|---|---|---|---|---|---|---|
| (Intercept) | -1.73 | [-3.48, 0.03] | | | | | | |
| youth_mean | 0.40** | [0.25, 0.55] | 0.09 | [0.06, 0.13] | .01 | [.00, .01] | .10** | |
| truth | 0.08 | [-0.08, 0.23] | 0.02 | [-0.02, 0.05] | .00 | [-.00, .00] | .05** | |
| | | | | | | | | $R^2$ = .010** 95% CI[.00,.02] |

Supplementary Table 12. Model 2 – Baseline, Parent-Report PDS
*Regression results using corrected_gap as the criterion*

| Predictor | *b* | *b* 95% CI [LL, UL] | *beta* | *beta* 95% CI [LL, UL] | $sr^2$ | $sr^2$ 95% CI [LL, UL] | *r* | Fit |
|---|---|---|---|---|---|---|---|---|
| (Intercept) | -1.43 | [-3.18, 0.32] | | | | | | |
| parent_mean | 0.43** | [0.29, 0.56] | 0.11 | [0.07, 0.14] | .01 | [.00, .02] | .11** | |
| truth | 0.05 | [-0.11, 0.20] | 0.01 | [-0.02, 0.05] | .00 | [-.00, .00] | .05** | |
| | | | | | | | | $R^2$ = .013** 95% CI[.01,.02] |

*Supplementary Figure 1.* Overlaid model predictions

A

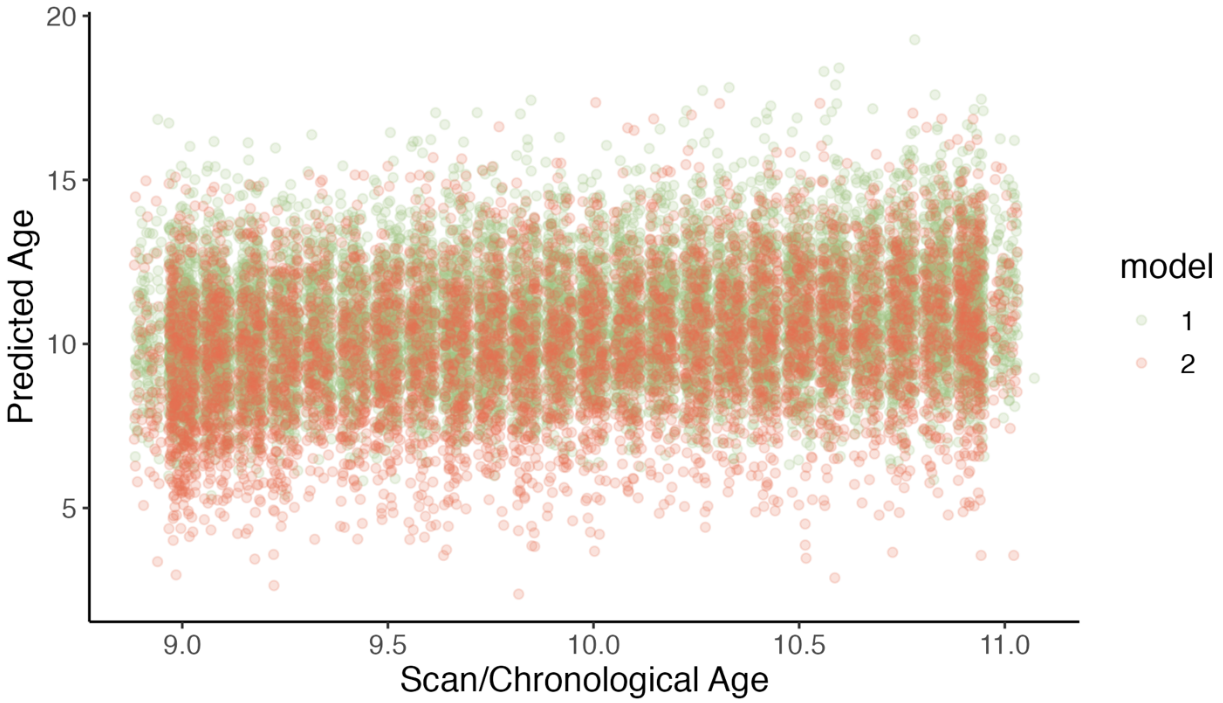

B

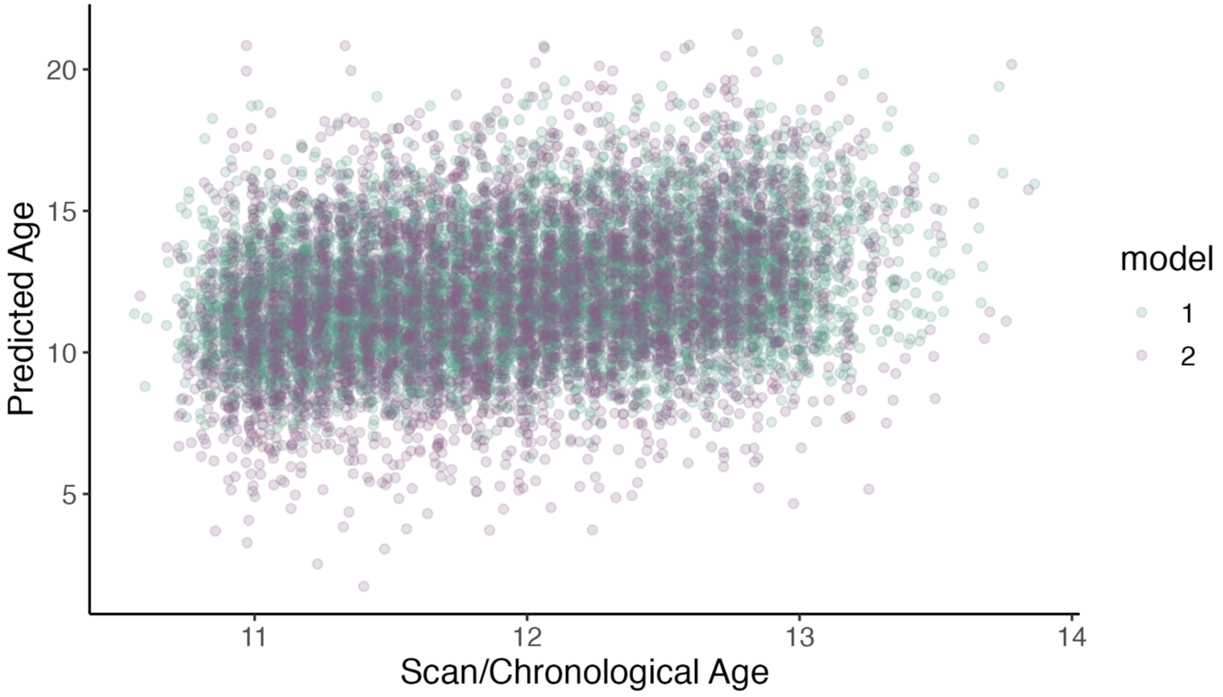

*Figure S1*. Overlaid model predictions. A) Both baseline models plotted on the same figure. B). Both follow-up models plotted on the same figure.
